# Induction of an immortalized songbird cell line allows for gene characterization and knockout by CRISPR-Cas9

**DOI:** 10.1101/2021.05.27.445896

**Authors:** Matthew T. Biegler, Olivier Fedrigo, Paul Collier, Jacquelyn Mountcastle, Bettina Haase, Hagen U. Tilgner, Erich D. Jarvis

## Abstract

The zebra finch is a powerful model for several biological fields, particularly neuroscience and vocal communication. However, this species lacks a robust cell line for molecular biology research and reagent optimization. Here we describe a cell line, CFS414, generated from zebra finch embryonic fibroblasts using the SV40 large and small T antigens. This cell line demonstrates an improvement over previous songbird cell lines through continuous and density-independent growth, allowing for indefinite culture and monoclonal line derivation. Cytogenetic, genomic, and transcriptomic profiling established the provenance of this cell line and identified the expression of genes relevant to ongoing songbird research. Using this cell line, we demonstrated a stress-dependent localization response of the zebra finch song nuclei specialized gene, SAP30L, and disrupted endogenous gene sequences using *S*.*aureus* Cas9. The utility of this cell line enhances the molecular potential of the zebra finch and validates cell immortalization strategies in a songbird species.

## INTRODUCTION

Songbirds have been utilized extensively to study vocal learning and communication (Brainard and Doupe, 2013; Petkov and Jarvis, 2012), sexual dimorphism (Choe and Jarvis, 2021; Wade and Arnold, 2004), comparative evolution (Chen et al., 2013; Fritz et al., 2014; Jarvis et al., 2013), and more. One songbird in particular, the zebra finch, has become an indispensable model organism for these diverse fields as it breeds easily in captivity (Mello, 2014). As a result, genomic resources (Korlach et al., 2017; Rhie et al., 2020; Warren et al., 2010), and neuroscience tools (Ahmadiantehrani and London, 2017; Garcia-Oscos et al., 2021; Heston and White, 2017; Long and Fee, 2008) have been more developed and characterized for the zebra finch than for any other songbird species (Mello, 2014). However, cell culture tools remain underdeveloped.

Cell culture is a valuable tool for model organisms to evaluate reagents, express recombinant genes, and characterize cellular and molecular processes in a controlled and quantifiable manner. Previously, zebra finch cell lines have been established using an induced pluripotent stem cell (iPSC) construct, STEMCCA, containing four transcription factors (Rosselló et al., 2013), or from tumors (G266 and ZFTMA) from a male and female, respectively (Itoh and Arnold, 2011). However, these lines are difficult to maintain as culture stocks compared to stable cell lines from other species and have not been widely utilized across the avian science community. Primary cells have also been utilized in several studies (Itoh et al., 2011; Kulak et al., 2020), though these cells have a limited lifetime *in vitro*. Primary cells also vary genetically between individuals and laboratories, risking issues with replicability between divergent zebra finch lab populations (Forstmeier et al., 2007). Particularly, none of these options have been reported to be capable of density-independent growth, which is useful for maintaining cell population homogeneity and also in generating transgenic cell lines.

The lack of a robust, stable cell line presents a key obstacle in furthering the introduction and optimization of molecular tools in the zebra finch, such as CRISPR-Cas9 (Ahmadiantehrani and London, 2017; Heston and White, 2017) Along with recent developments on transgenic strategies in the zebra finch (Gessara et al., 2021; Jung et al., 2019), an immortalized cell line would allow for the exploration of the basic molecular and cellular understanding of this vocal learning species. Isolating and resolving the particular role of key genes involved in vocal learning pathways is challenging without efficient cell lines to test their function and optimize reagents for *in vivo* experiments.

In poultry (*Galloanserans* e.g., chickens and ducks), cell lines have been successfully established. The DF-1 chicken cell line was derived from a spontaneously transformed culture population (Schaefer-Klein et al., 1998), while the DT40 chicken cell line was derived from a chicken infected with Rous sarcoma virus (Baba et al., 1985). Both of these lines have been critical for scientific and biotechnology applications (Cheng et al., 2018; Winding and Berchtold, 2001). Other avian cell lines include a duck embryonic fibroblast line, DEF-TA, that was immortalized with the Simian vacuolating virus 40 (SV40) large T antigen (Fu et al., 2012). Importantly, each of these cells are capable of robust, density-independent survival and proliferation, allowing for clonal propagation and antibiotic selection strategies. Such approaches have not been previously reported in passerine birds despite constituting over half (∼5,000) of all bird species (Ericson et al., 2003; Jarvis et al., 2014), representing a major gap in the toolset for studying this clade’s cellular and molecular biology.

Here, we established the provenance of an induced, continuous cell line from zebra finch embryonic fibroblasts with a lentivirus containing SV40 large and small T antigens (SV40Tt), capable of density-independent proliferation and clonal propagation. We characterized the line using Next Generation Sequencing (NGS) approaches, showing a stable cell line with a gene expression profile relevant to songbird research areas. We further demonstrated the cell line’s applicability in studying songbird genes related to vocal learning, and validated genome editing by CRISPR-Cas9 in the zebra finch genome.

## RESULTS

### Establishment of a continuous cell line derived from zebra finch

Building on studies of induced avian cell lines in poultry (Baba et al., 1985; Fu et al., 2012), we sought to generate an immortalized cell line from the zebra finch. We isolated finch embryonic fibroblast cells (FEFs) from Hamburger-Hamilton stage 28 embryos (Murray et al., 2013) (Figures 1A and 1B) and transduced them with lentivirus expressing either GFP (Figures S1A and S1B) or the SV40 large and small T antigen (SV40Tt, Figure 1C). Compared to GFP-transduced and wild-type FEFs, SV40Tt-transduced cells proliferated robustly beyond several passages (Figures 1D and 1E), outcompeting untransduced cell neighbors to become pure populations of immortalized cells. The most proliferative SV40Tt-transduced cell line was derived from a single continental chestnut-flanked white embryo, hereafter named CFS414 (RRID:CVCL_A7AT). The presence of SV40Tt in CFS414 genomic DNA was confirmed compared to primary FEFs (Figure 1F). Sextyping using the *CHD* gene (Griffiths et al., 1998) did not show the presence of the W chromosome, denoting a male embryo origin (Figure 1F).

**Figure 1.**
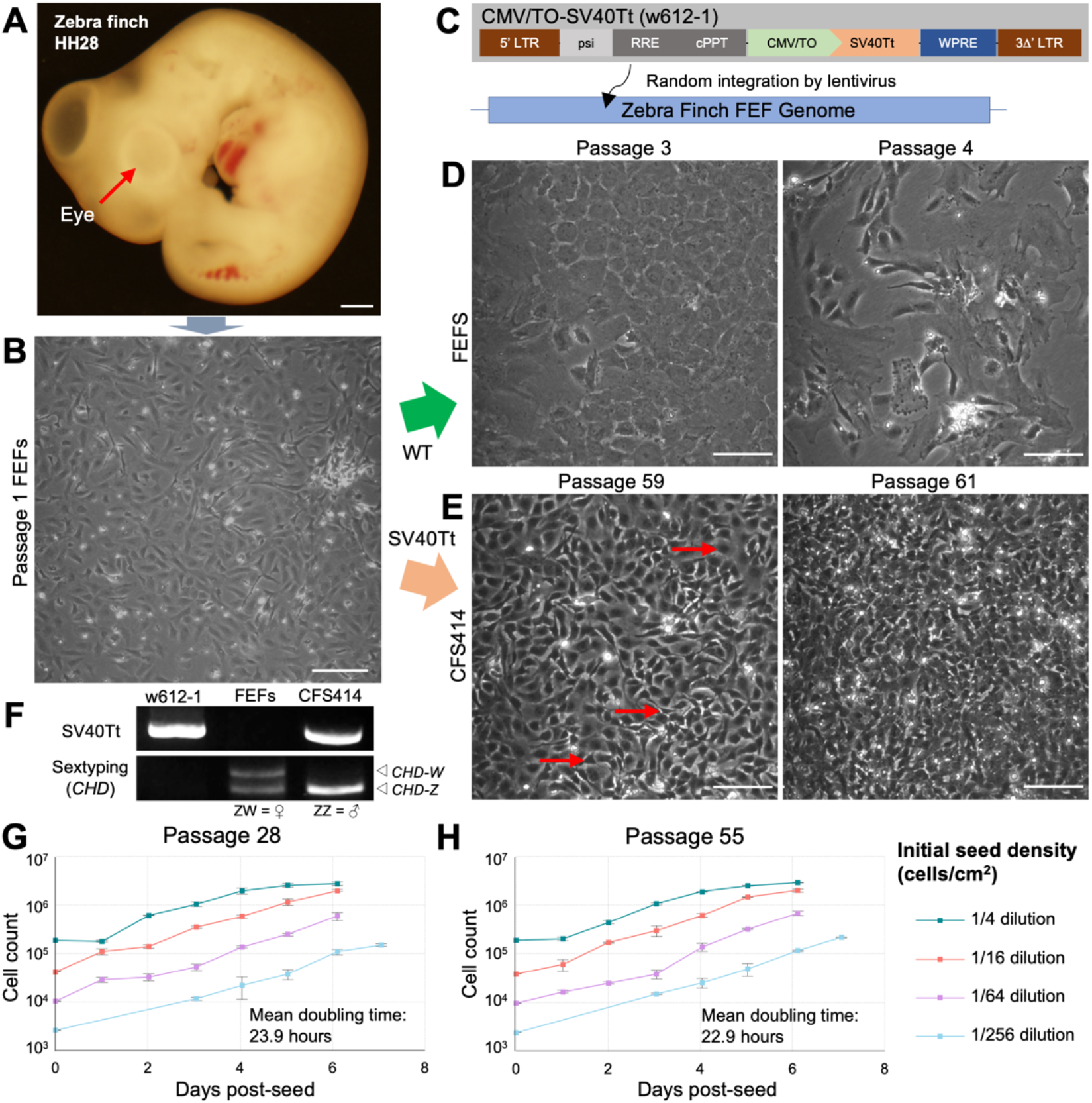
Generation of immortalized finch cells with SV40Tt. (A) Embryonic day 6 zebra finch with the continental chestnut-flanked white allele, imaged at 4x magnification. Note the lack of eye pigmentation (red arrow) for this color-coat allele. Scale bar = 200µm. (B) Exemplar image of passage 1 finch embryonic fibroblast (FEF) culture. (C) Diagram of lentiviral construct (w612-1) delivering SV40Tt immortalization factor into the zebra finch genome. Abbreviations: LTR 5’ and 3Δ’, Long terminal repeat sites; psi, retroviral packaging element; RRE, Rev responsive element; cPPT, central polypurine tract; CMV/TO, Cytomegalovirus promoter with 2 Tet responsive element motifs, allowing for inducible expression in the presence of Tet-repressor constructs; SV40Tt, Simian virus 40 large and small T antigen; WPRE, woodchuck hepatitis virus posttranslational responsive element. (D) Passage 3 FEFs at full confluence (left) and passage 4 from a 1:2 split (right), with low confluence 11 days after seeding. For further FEF characterization, see Figure S1. (E) CFS414 cells between passage 59 at full confluence (left) and passage 61 from a 1:10 split (right), overconfluent 5 days after seeding. Note the presence of multiple, distinct cell morphologies (red arrows). (F) PCR amplicons of SV40Tt and sex-chromosome specific *CHD* gene sequence (ZZ is male, ZW is female), using w612-1 and genomic DNA derived from primary FEFs and CFS414 cells (right). (G-H) Growth curves of CFS414 cells at (G) passage 28 and (H) passage 55. Seeding densities range from 1:4 (160,000 cells/well) and 1:256 (1,250 cells/well) dilution, each showing a log-phase doubling time of approximately 23.9 hours. Doubling time between passage 28 and passage 55 were similar (p=0.553, two-tailed Student’s t-test). Error bars denote SEM.

Compared to wild-type FEF controls (Figure S1C), CFS414 cells were highly proliferative (Figures 1G and 1H). These cells also grew independently of initial seed density, maintaining an average doubling time of 23.9 hours during log-phase growth (Figure 1G), similar to other cell lines such as HEK293T cells (RRID: CVCL_0063; doubling time ∼24-30 hours). At low plating densities, CFS414 cells are able to propagate and form colonies from individual cells, making monoclonal selection possible. At high confluence, CFS414 cells continue growing with limited contact inhibition, and do not form “collagenated sheets” due to overgrowth, as seen in primary FEFs (Figure S1D and S1E). The log-phase doubling time remained stable even after another ∼30 passages (Figure 1H), suggesting low population-wide senescence.

### Assessment of chromosome and genome sequence integrity

Chromosomal abnormalities are common among cell lines (Macville et al., 1999), particularly those transformed by SV40 T antigens (Ray et al., 1992), and may impact the generalizable nature of cell lines to a species’ biology. G-banded karyotypes of macrochromosomes were performed on CFS414 cells. At passage 38, most chromosomes appeared normal and intact compared to previous studies (Itoh et al., 2011; da Silva dos Santos et al., 2017; Rosselló et al., 2013), but trisomies in many chromosome pairs were common (Figures 2A and 2B; Figures S2A and S2B) and varied across cell spreads within the sampled populations. Passage 63 revealed a more variable karyotype (Figures S2C and S2D). However, a monoclonal line derived through flow cytometry (denoted clone F6) showed a similarly aberrant but relatively less variable karyotype (Figures 2C and 2D; Figures S2E and S2F). In all analyses, multiple copies of the Z chromosome were indicated, as well as a what appeared to be a mutant Z chromosome variant (Z’; Figures 2C and 2D).

**Figure 2.**
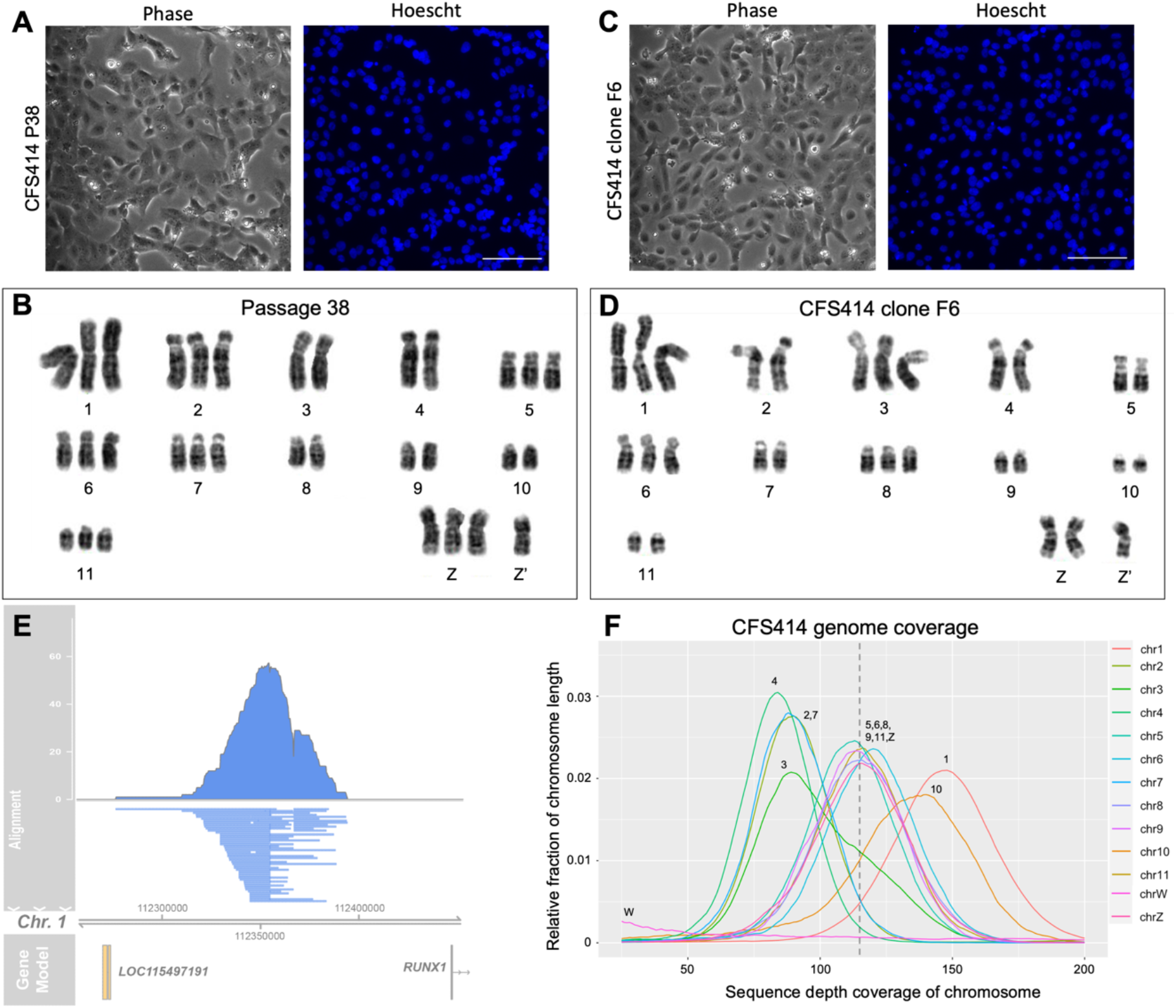
Variability of CFS414 cells is not defined by polyclonal SV40Tt integration. (A, C) Phase contrast images and accompanying Hoescht33342 nuclear stain images of (A) passage 38 and (C) CFS414 clone F6 cells derived by flow cytometry. (B, D) Corresponding Giemsa-stained macrochromosomes for (B) Passage 38 and (D) F6 cells, sorted by chromosome pair. Trisomies are present for many chromosomes, with variability across cells. Chromosomes are arranged and numbered by size, with linkage-mapped chromosome numbers from previous studies listed in parentheses. Note presence of mutant Z chromosome (Z’) in both karyotypes. See Figure S2 for further karyotype spreads. (E) Pileup of mapped Pacbio subreads containing the SV40Tt provirus from CFS414 cells, showing SV40Tt insertion in the intergenic space between *RUNX1* and *LOC115497191* on chromosome 1. (F) Relative coverage for each chromosome in CFS414 cells. Three distributions of peaks in the cell line coverage plot suggests cell population-wide copy number differences between chromosomes. The dotted line represents the expected coverage (115X) and the relative average copy number for all chromosomes. See Figure S3A for individual chromosome coverage plots.

To determine whether these variable populations represented multiple transduction events within a polyclonal population or from a single event, the genome was sequenced at 115X coverage on passage 31 CFS414 cells (Table S1). Pacbio subreads were filtered using the Basic Alignment Search Tool (BLAST) (Altschul et al., 1990) to identify subreads containing the SV40Tt proviral sequence. The selected reads were mapped to the high-quality zebra finch reference genome assemblies, generated by the Vertebrate Genomes Project (VGP) (Korlach et al., 2017; Rhie et al., 2021), to identify the site of integration (Figure 2E). Proviral sequences were found in 115 reads, and 111 mapped to a single insertion site at chromosome 1, in the intergenic space between the *RUNX1* gene and the uncharacterized *LOC115497191* gene. The other 4 provirus-containing subreads did not align to the reference genome, likely owing to low subread sequence quality. These data suggest that, by passage 31, CFS414 cells originated from a single cell transduced with SV40Tt.

We then aligned the Pacbio reads to the reference genome to calculate the coverage in each chromosome. Sequence coverage varied between chromosomes, indicating differences in population-level copy number between chromosomal pairs, as seen in the karyotypes (Figure 2D; Figure S3A). Certain chromosomes appeared to group together in coverage, with most peaking around the expected coverage (115X). Others showed significantly higher or lower sequence depths (85X and 140X), these peaks representing relative proportional copy number differences (74% and 122%, respectively) between chromosomes across the cell line population, consistent with the variably polyploidy seen in the karyotypes. By comparison, differences in coverage by chromosome pair were not seen in the reference genomes, except in the female sex chromosomes (Figures S3B-S3E). Finally, the Pacbio reads were assembled to produce a high-quality contig assembly for the CFS414 genome. The assembly length of the primary pseudo-haplotype was 1.05Gb, close to the expected 1.1Gb (for sequence assembly statistics, see Table S1).

Structural variants were determined using Bionano optical mapping data. The raw Bionano reads were assembled into a physical map and aligned to the male reference genome (Korlach et al., 2017; Rhie et al., 2021). From this alignment, we identified several hundred large (≥150 kb) structural variants (indels, inversions, copy number variations, and translocations) in the CFS414 genome (Figure S3F). The most significant variant we found was near one end of the Z chromosome (Figure S3F), denoting several large changes possibly related to the polymorphic Z chromosome identified in the karyotypes. Across the genome, most variants we found were copy number variations, consistent with the observed aneuploidy. For the other structural variations, it could not be determined which originated naturally or were the result of SV40Tt mutagenesis. We note, however, that such variations are typical between ‘wildtype’ genomes, as recent VGP genome assemblies have shown a higher rate than previously expected (affecting ∼1.3% of the genome) between haplotypes or individuals (Rhie et al., 2021; Yang et al., 2021).

### Single-cell RNA sequencing reveals CFS414 transcriptomic landscape

We observed that the CFS414 cell line contains several distinct morphological populations varying in size, optical thickness, packing density, and shape (Figure 1E, red arrows), and these morphological characteristics persisted in the F6 monoclonal line (Figure 2B). We wondered whether these morphologies indicated transcriptomic instability or consistent differentiation into different cell types.

To catalog the transcriptomic landscape of CFS414 cells, we used single-cell RNA sequencing (scRNAseq) for samples at passage 24 and 50 (Figures S4A-S4D). In all, 13,108 cells were sequenced (for sequence statistics, see Tables S2 and S3). From a total of 22,139 annotated genes, transcripts for 15,719 genes were identified in at least 0.1% of cells (Table 1; Table S4), representing a sizeable portion of zebra finch genes. Curated gene lists were used to determine the expression of genes pertinent to avian specialized traits or disease research. The expression of 19 genes associated with West Nile Virus infection in zebra finch (Newhouse et al., 2017) and 19 genes with Avian Influenza infection in chicken (Drobik-Czwarno et al., 2018) were found (Table 1; Table S4), highlighting the ability to study viral infections. Of 5,472 genes with specialized regulation in the zebra finch vocal learning brain nuclei compared to surrounding regions (Gedman et al., in preparation), 2,967 genes were expressed in more than 10% of the cells (Table 1; Table S4). CFS414 cells also significantly expressed 908 identified gene orthologs associated with neurological speech disorders, taken from the Human Phenotype Ontology database (Köhler et al., 2020).

**Table 1.** Expression of gene lists related to different research fields utilizing zebra finches. See Table S4 for further expression statistics. Abbreviations: WNV, West Nile Virus; DEG, differentially expressed gene; HPO, Human Phenotype Ontology.

| **Genelist (# identified genes)** | **# Expressed**<br>**≥0.1% cells** | **# Expressed**<br>**≥10% cells** |
| --- | --- | --- |
| **Zebra Finch Genes (22139)** | 15719 | 9915 |
| **WNV Genes (25)** | 19 | 8 |
| **Avian Influenza Genes (21)** | 19 | 15 |
| **Song Nuclei DEGs (5472)** | 5485 | 2967 |
| *Upregulated genes (3129)* | 2669 | 1744 |
| *Downregulated genes (2627)* | 2089 | 1323 |
| **HPO disordered speech genes (1184)** | 1090 | 908 |

We next assessed the transcriptomic stability of the cell line. Averaging the gene expression for each passage, we found that their respective gene expression profiles were highly concordant, with only 195 genes differing by a natural log fold change (logFC) cutoff of more than 0.25 (Figure 3A; Table S4). The majority of these genes (n=126) decreased in expression, and the cell-by-cell variability of the 200 most variable genes also went down between passages (Figure 3B; Figure S3E). PCA-mediated cell clustering of a combined passage dataset (Butler et al., 2018; Hafemeister and Satija, 2019) resolved 5 clusters across both passages, visualized by uniform manifold approximation and projection (UMAP) (Figure 3C and 3D; Figures S4E and S4F). Most clusters were closely packed toward the center of the UMAP, with few sharp boundaries occurring between clusters. The top expression markers between several clusters included many ribosomal and cell cycle associated genes (Table S5), despite regression analysis of the latter group prior to cluster identification. Indeed, cell cycle analysis confirmed a high proliferation rate varying across clusters, with more than 30% of cells in the G2M phase of both passages (Figure S4G). We found that many of the most variable genes decreasing between passages 24 and 50, such as *TNNT2* (Figure 3E), were related to passage 50 reductions in the rightmost portion of cluster 2 compared to passage 24 (Figure 3C and 3D, black arrows). This difference in cluster demographics (Figure S4F) between passages is most likely due to progressive passaging modifications or slight differences in cell confluence at the time of sequencing.

**Figure 3.**
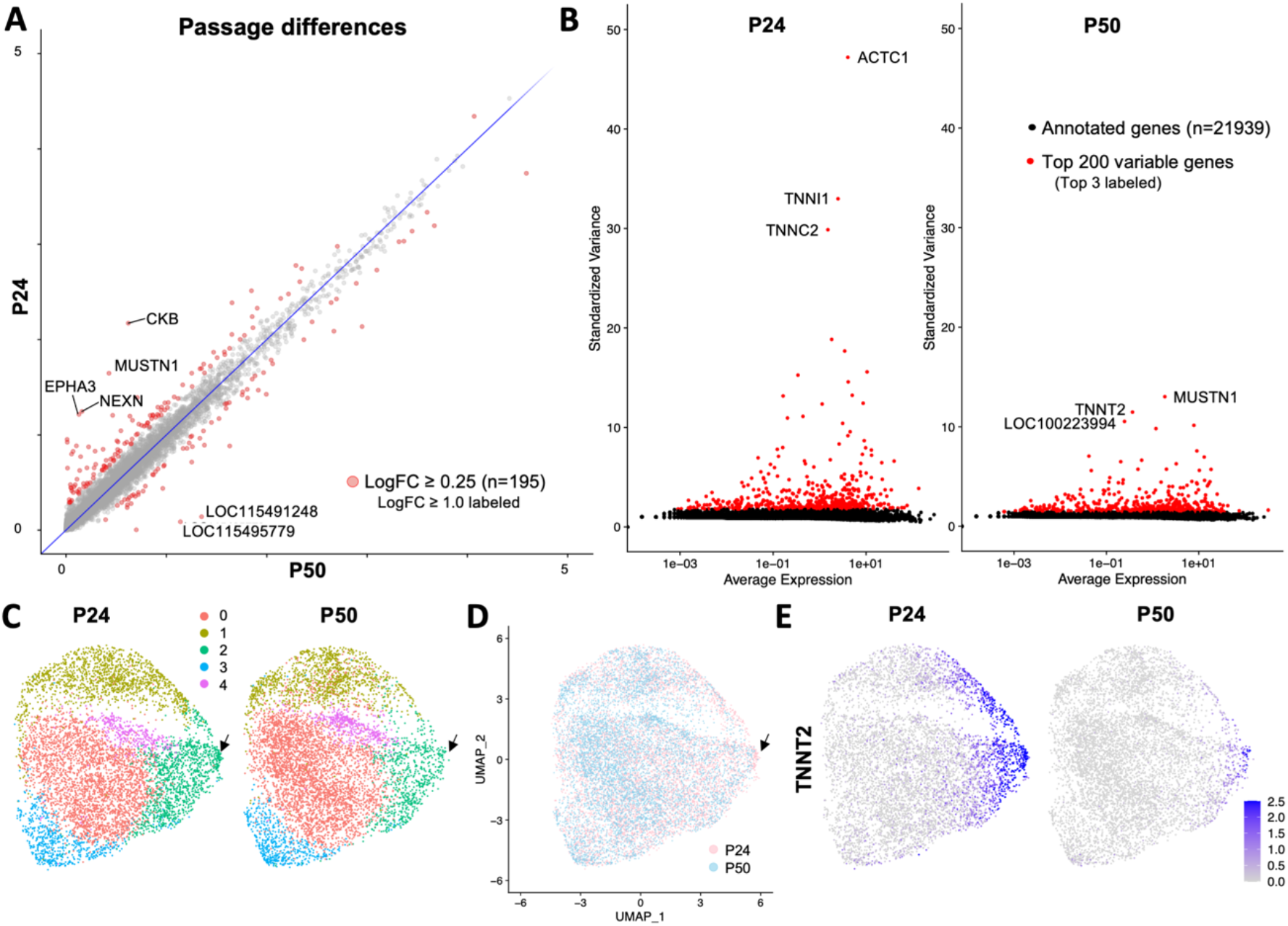
Single-cell RNA sequencing reveals relative homogeneity of CFS414 cells. (A) Average expression of genes in passage 24 (P24, y-axis) and passage 50 (P50, x-axis) cell samples. Red markers highlight a natural log fold change (logFC) ≥ 0.25; genes with logFC ≥ 1 are labeled. (B) Plot of all annotated genes by standardized variance (y-axis) and average expression (x-axis) for P24 (left) and P50 (right). Note the difference in the relative decrease in variance of the top 200 genes (red) between passages. See Figure S4D for these plots at a y-axis scale of 0 to 5. (C) UMAP plots of integrated P24 (left) and P50 (right) samples, with clusters labeled by color. (D) Overlay of passage 24 (pink) and passage 50 (blue) UMAP plots. Black arrows highlight passage difference in expression in cluster 3. (E) UMAP plot colored for *TNNT2* expression between passages.

To determine what cell type CFS414 cells most resembled, we looked at the gene ontology (GO) terms for differentially expressed genes between cell clusters. Cluster 2 was the most distinct cluster (Figure S4E), with a GO gene profile corresponding to muscle function (Table S6). In particular, the myocyte differentiation markers *MYOG* and *MEF2C* were elevated, as well as several cardiac myocyte-associated troponins and actins (Figure 4A; Table S5). Across all clusters, several gene markers for muscle cell types (Bryson-Richardson and Currie, 2008) (e.g., *MYOD1, SIX1*) were also present (Figure 4B). GO Terms for several other clusters were also related to muscle cells, including muscle contraction and actin-myosin filament sliding (Table S6). These findings suggest that the originally transduced CFS414 cell was derived from myogenic tissue.

**Figure 4.**
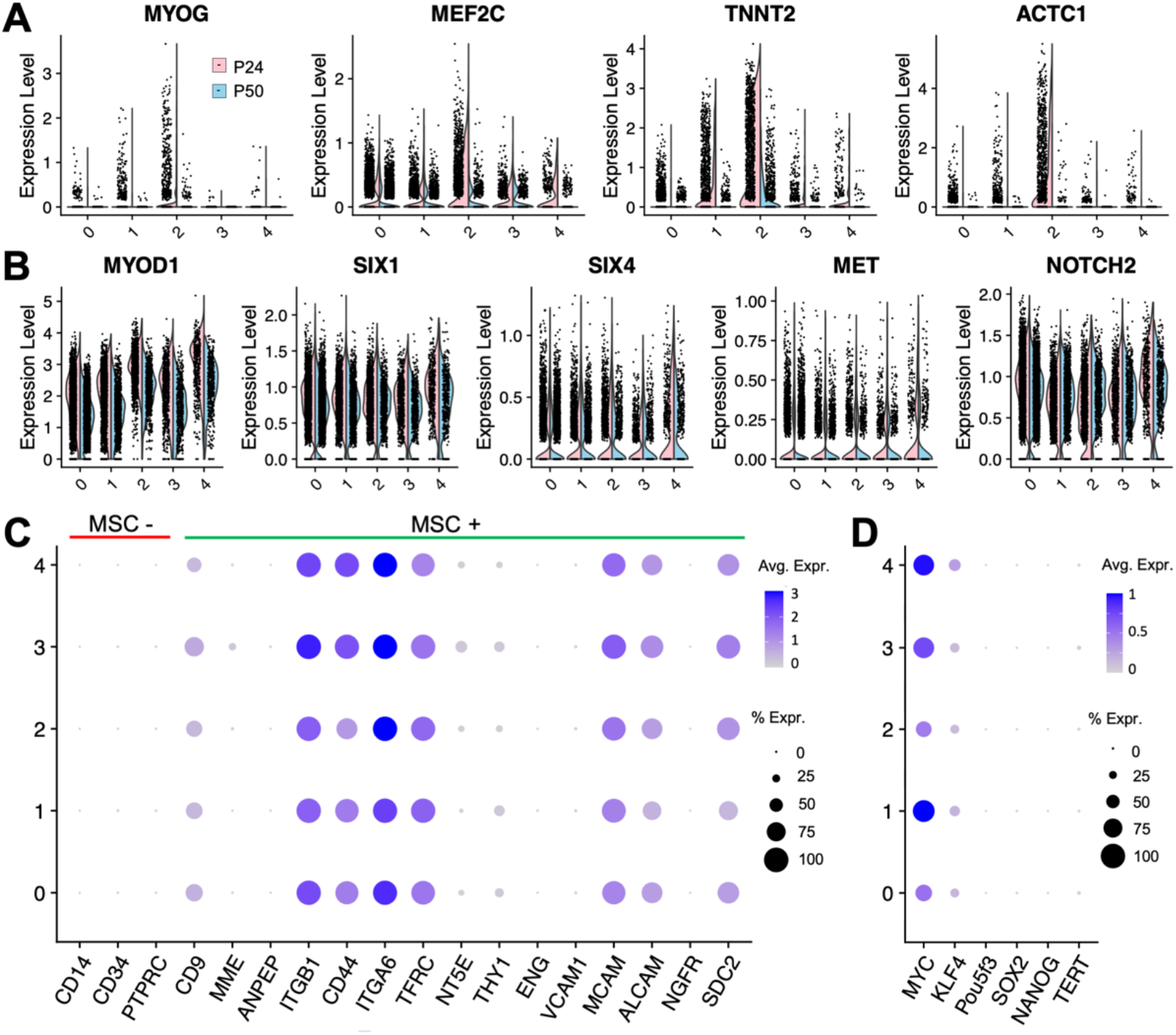
Myogenic cell identity of CFS414 cells. (A-B) Violin plots of (A) cardiac myocyte gene markers and (B) muscle cell gene markers in CFS414 cells, split by passage. (C) Dot plot of negative (red line) and positive (green line) mesenchymal stem cell (MSC) markers in passage 24 CFS414 cells. (D) Dot plot of canonical pluripotency markers in passage 24 CFS414 cells.

CFS414 cells also expressed stem cell markers. In all clusters, significant expression was seen for several markers associated with mesenchymal stem cells (Fitzsimmons et al., 2018) (Figure 4C). The pluripotent gene markers *KLF4* and *MYC* were also highly expressed, but not *SOX2* or the avian *OCT4* ortholog, *Pou5f3* (Figure 4D). Expression of *TERT* and *NANOG* were also very low. Evidence for some state of cell potency was also apparent based on the varied morphology appearing in culture (Figure 1E, red arrows).

### Stress-responsive SAP30L localization in the zebra finch

Cell lines are powerful tools for molecular and cell biology, and the CFS414 line represents an ability to robustly explore gene roles in songbirds. To validate the cell line’s applicability for studying zebra finch gene function, we manipulated expression of one gene upregulated in HVC and other song learning nuclei relative to the surrounding brain subdivisions, *SAP30L* (Lovell et al., 2020; Olson et al., 2015) (Figures 5A and 5B). The HVC song nucleus controls the timing and sequencing of learned song (Long and Fee, 2008). *SAP30L* has been extensively studied in mammalian cells, but not avian species. SAP30L and the SAP30 paralog interchangeably form part of the histone deacetylation (HDAC) complex (Laherty et al., 1998; Viiri et al., 2006; Zhang et al., 1998). These proteins engage in non-specific binding to DNA, bending it to increase adjacent histone proximity to the protein complex and enhance HDAC-mediated deacetylation of histone tails in larger genomic areas, repressing gene transcription (Viiri et al., 2009). SAP30L contains a binding domain for phosphatidylinositol monophosphates (PIPs) that overlaps the nuclear localization signal (NLS), inducing SAP30L expulsion from the nucleus upon binding. In mammals, nuclear PIP levels are known to increase following oxidative stress (Shah et al., 2013), making SAP30L a redox-dependent transcriptional de-repressor. In the zebra finch, the vocal learning behavior requires high-frequency excitatory firing through NMDA receptor activation (Ding and Perkel, 2004). Such firing properties have been shown in mice to induce excitotoxicity and expansive oxidative stress (Reyes et al., 2012). Genes upregulated for neuroprotection in song nuclei have been proposed to protect the cells from senescence due to the increased activity compared to the surrounding brain subdivisions (Hara et al., 2012; Horita et al., 2010).

**Figure 5.**
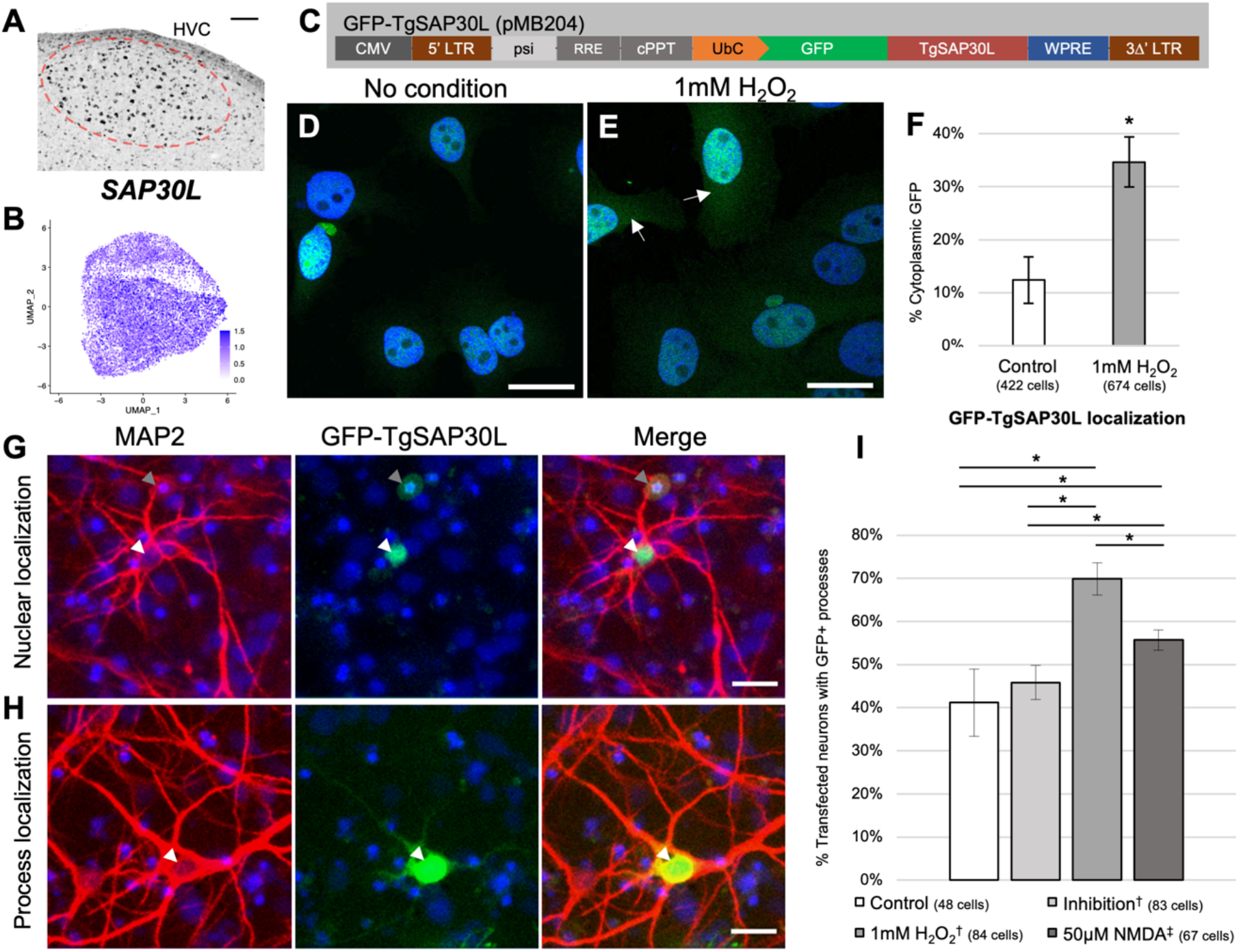
SAP30L localization in zebra finch cells is affected by oxidative stress. (A) SAP30L mRNA detected by *in situ* hybridization in the zebra finch HVC song nucleus (red circle), taken from the Zebra finch Expression Brain Atlas (ZEBrA) (RRID:SCR_012988). Scale bar = 100*µ*m.(B) UMAP plot showing SAP30L expression in CFS414 cells (passages combined). (C) Illustration of the pMB204 GFP-SAP30L lentiviral construct. (D-E) DAPI labeled CFS414-pMB204 cells under (D) control or (E) H2O2-mediated oxidative stress conditions. Arrows denote cytoplasmic localization. (F) Percent of CFS414-pMB204 cells showing cytoplasmic GFP signal in H2O2 and control media conditions. (G-H) MAP2 (red) and DAPI (blue) labeled primary cultured neurons transfected with pMB204 plasmid (white arrowheads) showing (G) nuclear or (H) process localization of GFP-SAP30L (green). Grey arrowheads denote autofluorescence. (I) Assessment of GFP localization in control media conditions, ion channel inhibition (†, for inhibitors, see Table S7), or challenged with either 1mM H2O2 or 50µM NMDA (‡, Table S7). * denotes significance, p < 0.05 based on (F) two-tailed Student’s t-test or (I) one-factor ANOVA. Error bars denote SEM. Scale bars = 20µm.

To assess whether SAP30L has a stress-dependent translocation response in the zebra finch, *in vitro*, CFS414 cells were transduced with lentivirus containing GFP fused to the zebra finch SAP30L protein coding sequence (GFP-TgSAP30L, Figure 5C). At baseline controls, these cells demonstrated strong nuclear and nucleolar GFP signal (Figure 5D). These cells were then challenged with an H_2_O_2_-mediated assay of oxidative stress previously used in human cell lines (Viiri et al., 2009). In response, they subsequently showed a 3-fold increase in cytoplasmic GFP signal compared to control cells (Figures 5D-5F), indicating that the redox-dependent shuttling behavior of SAP30L is conserved between avian and mammalian cells.

To substantiate the CFS414 localization experiments in primary cells and to investigate the unstudied role of SAP30L in the nervous system, we examined whether SAP30L shuttling behavior occurs in neurons. Zebra finch primary cortical neurons were prepared and transfected with plasmid containing the same GFP-TgSAP30L construct. Transfected primary neurons showed differential localization on a cell-by-cell basis (Figures 5G and 5H). In H_2_O_2_-challenged neurons, the percent of cytoplasmic GFP signal increased (including in the MAP2-labeled neuronal processes) compared to control or ion channel inhibited neurons (Figure 5I). NMDA-mediated excitotoxicity (Zhou et al., 2013) also caused an increase in cytoplasmic GFP signal, though not as high as H_2_O_2_.

### Knockout of zebra finch genes in CFS414 cells using CRISPR-Cas9

The use of genome editing tools such as CRISPR-Cas9 has not been previously reported in songbirds, in part due to the lack of robust genetic substrates for testing and optimizing these tools. To assess editing efficiency in the zebra finch genome using CFS414 cells, we generated a two-plasmid system for CRISPR-mediated gene ablation. One plasmid is a *piggyBac* transposon vector (pMB952) with a flip-excision (FLEx) cassette that expresses either nuclear GFP or *S*.*aureus* Cas9 (SaCas9; Figures 6A and 6B). The other is a plasmid to be used in tandem with a Tol2 transposon vector (pMB1052), containing a U6-driven guide RNA (gRNA) cassette and Cre recombinase to activate SaCas9 expression (Figure 6C). These vectors are selectable using puromycin or neomycin, respectively. This plasmid system was then used to deliver a GFP-targeting gRNA, which was then transfected with their respective transposase vectors into a monoclonal line expressing GFP (clone 2D4) (Figures S5A-S5E) and purified by antibiotic selection. We observed a significant decrease in GFP signal compared to a control gRNA targeting the bacterial *LacZ* gene (Figure S5F).

**Figure 6.**
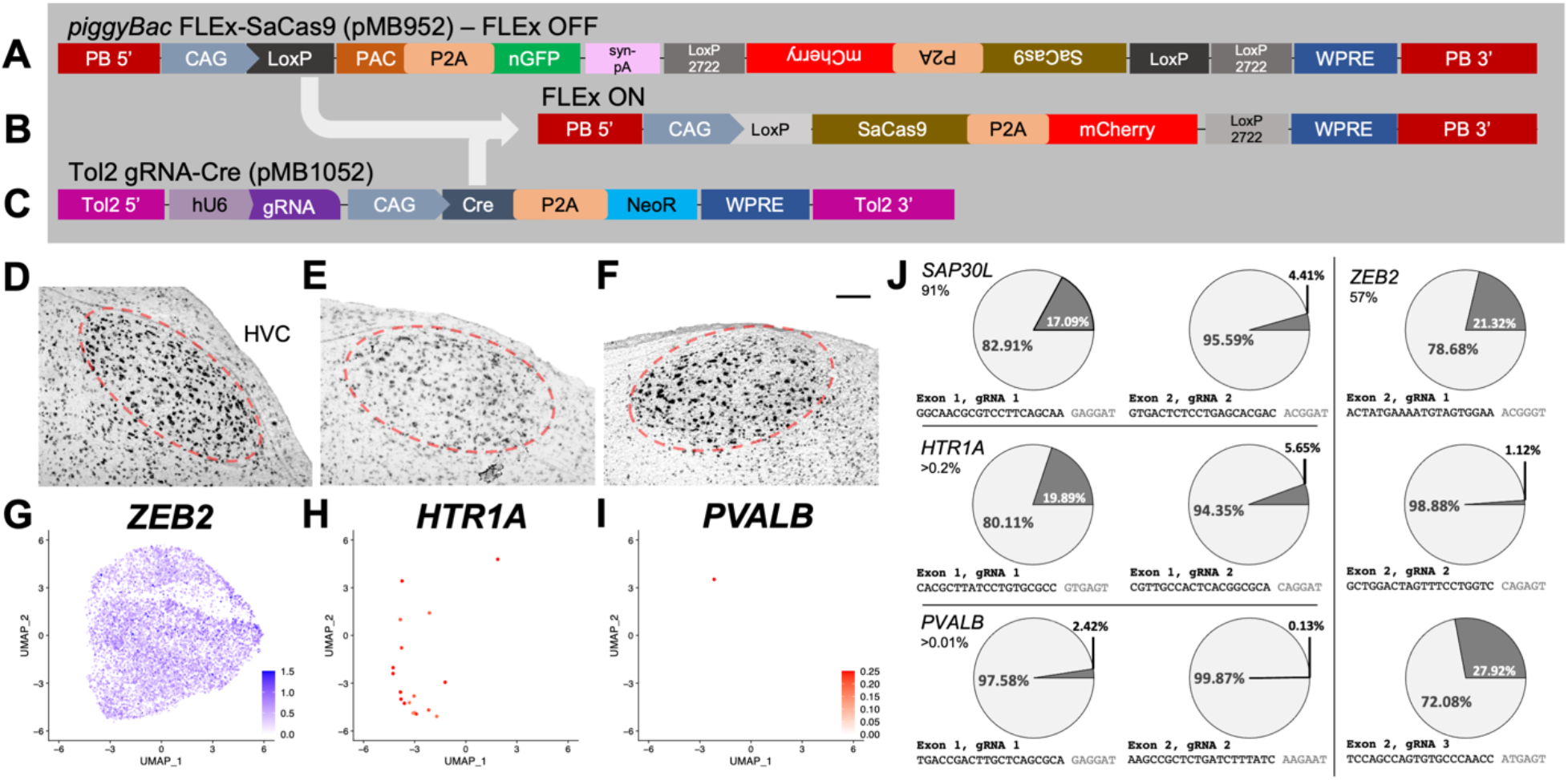
Indel formation in the CFS414 genome using SaCas9. (A) The pMB952 *piggyBac* transposon plasmid is a FLEx-cassette vector for Cre-conditional expression. Under constitutive expression, the CAG promoter expresses PAC puromycin resistance gene, with a P2A cleavage peptide sequence and nuclear GFP (nGFP). (B) In the presence of Cre, the PAC-nGFP sequence is excised and SaCas9-P2A-mCherry is flipped for sense expression under the CAG promoter. (C) The pMB1052 *Tol2* transposon plasmid is for Cre-mediated expression of SaCas9 in pMB952-integrated cells, with the delivery of a DNA-targeting gRNA. (D-F) Chromogenic *in situ* hybridization of (F) *ZEB2*, (G) *HTR1A*, and (H) *PVALB* mRNA expression in the zebra finch HVC song nucleus (red circle). Imaged at 10x magnification, scale bars = 100µm. (G-I) UMAP plots colored for (C) *ZEB2* (D) *HTR1A*, and (E) *PVALB* expression in CFS414 cells (combined passages). Note the difference in scales for genes expressed at high (blue) and low (red) levels. (J) Pie charts showing the percentage of modified genome loci at a gRNA target site from CFS414-pMB952 clone 1G10 cells transfected with gRNAs targeting endogenous zebra finch genes. Each targeted gene (percent of CFS414 cell expression provided below gene name) is separated by lines and gRNA sequence and protospacer adjacent motif (PAM) is listed below respective pie chart. For specific indel sequences, see Figure S6. Abbreviations: PB 5’ and 3’, *piggyBac* transposon sites; bGH pA, bovine growth hormone polyadenylation site; Syn pA, synthetic polyadenylation site.

To assess whether Cas9 would target endogenous zebra finch genes in CFS414 cells, we generated a monoclonal cell line expressing the FLEx-Cas9 vector, denoted as CFS414-pMB952 clone 1G10 (Figures S6H-S6K). We then transfected this cell line with Tol2 gRNA-Cre vectors targeting *SAP30L, ZEB2, HTR1A*, and *PVALB*. These genes are each upregulated in the HVC song nucleus (Hara et al., 2012; Lovell et al., 2020; Pfenning et al., 2014) (confirmed here in Figure 5A and Figures 6D-6F) and differ greatly in their relative expression in CFS414 cells (Figure 5B; Figures 6G-6I). Neomycin selection and subsequent purification by fluorescence-activated cell sorting (FACS) increased the relative population of SaCas9-mCherry-positive cells (Figures S6I-S6J). Using an established method for indel detection (Nelson et al., 2015), deep sequencing of PCR-amplified genomic loci surrounding the gRNA target demonstrated variable cutting efficiencies in 1G10 cells, with indel formations found in 0.1-27% of sequenced amplicon reads, depending on the gRNA used (Figure 6J). Indels ranged in size from -83 to +106 bp, though the majority of detected indels fell within 15 bp (Figure S6A-S6M). Predicted off-target sites for 4 of the most efficient gRNAs showed very low indel formation rates, similar to untargeted control amplicons (Figure S6N). The gRNAs targeting *ZEB2* were the most effective, and those for *PVALB* gene were the least effective; this may be the result of intrinsic gRNA sequence or cell line properties, such as poor chromatin accessibility at the target site related to non-expression (Jensen et al., 2017; Figure 6F). Overall, these targeted indel formations provide evidence that CRISPR-Cas9 may be used to induce frameshifts in endogenous genes and disrupt protein expression in the zebra finch.

## DISCUSSION

CFS414 cells overcome the obstacles of previous cell lines, fulfilling an unmet need for a robust and readily manipulable zebra finch cell line. These cells represent the first published instance of an induced cell line in any songbird species, providing insight on the cell biology of a diverse and distinct avian clade. The previous G266 and ZFTMA zebra finch cell lines (Itoh and Arnold, 2011) have been instrumental in the study of mRNA expression (Balakrishnan et al., 2012), antibody validation (Condro and White, 2014), microRNA characterization (Lin et al., 2014a), and DNA methylation (Steyaert et al., 2016), highlighting the importance of a zebra finch cell line. However, we and others (Carlos Lois, personal communication) have noted difficulty in maintaining these cell lines in the lab due to slow growth, low cryogenic viability, and requirements for a high seeding density. These previous cell lines are also not currently available through commercial or non-profit cell repositories. Such characteristics can limit a cell line’s usefulness and application, hindering their widespread adoption by the research community. The CFS414 and derivative cell lines will be maintained and distributed by the Rockefeller University Antibody & Bioresource Core Facility.

The robust and versatile nature of the homogeneous CFS414 cell line enables strategies to generalize protocols and replicate findings between labs. The identification of a single SV40Tt integration point supports the origin of CFS414 cells from a single cell, likely representing the most proliferative of the transduced cells that displaced other populations across passage dilutions. Critically, density-independent proliferation is possible with this cell line, critical for monoclonal and transgenic cell line generation to improve and maintain population homogeneity.

The CFS414 genome assembly and karyotypes showed aneuploidy. SV40Tt has been shown to inhibit p53, (Lane and Crawford, 1979; Ray et al., 1992) a tumor suppressor involved in the repair of DNA damage (Kastenhuber and Lowe, 2017) which could lead to chromosomal abnormalities. The different chromosomal groupings by sequence depth coverage provides a sense their proportional, population-level copy number in CFS414 cells, further shown with the cytogenetic and structural variant analyses. Continuous cell lines, including those generated with SV40Tt, are known to be associated with some chromosome rearrangements. Indeed, significant aneuploidy is common among the most utilized cell lines, such as HEK293T (Lin et al., 2014b), HeLa (Macville et al., 1999), and DT40 cells (Chang and Delany, 2004). The dynamics of SV40Tt expressed in CFS414 cells have not been directly characterized, though genome and scRNAseq analysis revealed a closely related population with minor variation. Nonetheless, it is generally recommended staying below a certain number of passages in cell line culture before restarting with initial frozen stocks (Hughes et al., 2007; Masters and Stacey, 2007; Vunjak-Novakovic and Freshney, 2006), and we believe proper cell culture practices will mitigate any deleterious effects on CFS414 cell line applications.

Originating from a male embryo, CFS414 male cells are useful in the context of vocal learning research, as singing behavior in zebra finches is restricted to males. Transcriptomic analysis determined gene expression profiles suitable for the study of many genes important to vocal learning research, as well as other fields, such as avian virology. The presence of a distinct Z chromosome variant is interesting, as Z chromosomes polymorphisms have been found in zebra finch laboratory populations (Itoh et al., 2011). While it is unknown whether this structural variation originated from the embryo or from SV40Tt mutagenesis, we note that the original embryo presented the continental chestnut-flanked white allele, which characteristically lacks eye pigmentation across development (Figure 1A). This allele is known to be sex-linked in zebra finches (Landry, 1997), though the genetic mechanism has not been precisely determined. Further investigations with this cell line and animal populations may determine whether the coat-color allele and the variant Z chromosome region are related. While this cell line is not representative of the entire zebra finch genetic landscape, the demonstration of this immortalization technique may be applied to generate cell lines from other tissue samples to study chromosomal polymorphisms, the female W chromosome, or germline restricted chromosome (Kinsella et al., 2019; Pigozzi and Solari, 1998).

The CFS414 cell line expressed several gene markers for human myoblast cells, with morphological and transcriptomic evidence of active differentiation observed. The additional expression of a few pluripotent gene markers suggests that CFS414 cells exist in some potent or progenitor cell state. Previous studies have demonstrated some limited potency of myoblast cells (Nihashi et al., 2019; Watanabe et al., 2004), and other studies have shown that SV40 T antigen expression can drive the expression of potency markers (Batsché et al., 1994). Future work can explore protocols to determine and optimize guided differentiation of CFS414-derived populations, potentially halting myogenic differentiation or differentiating other cell types, such as neurons (Watanabe et al., 2004; Xiao et al., 2018).

Cell lines such as CFS414 allow for the testing and optimization of tools and reagents, such as viral vectors or antibodies, before utilizing them *in vivo*. Overexpression of zebra finch genes can help identify their functional role within the context of its natural cell environment. Our SAP30L case study demonstrated redox-dependent shuttling into the cytoplasm, as previously documented in human cells (Viiri et al., 2009). Our results further showed the novel finding of SAP30L translocation following NMDA receptor activation.

As SAP30L is differentially upregulated in several zebra finch song nuclei (Lovell et al., 2020; Olson et al., 2015), these data imply SAP30L cytoplasmic translocation could play a specialized role in vocal learning through activity-dependent gene de-repression following singing behavior. As our tested stimuli for the SAP30L migration assays are sources of oxidative (H_2_O_2_) and excitotoxic (NMDA) cellular stress, SAP30L may also have a potential neuroprotective role in the highly active zebra finch song system. Future studies to conclusively determine these roles *in vivo* would be supported by reagent validation in CFS414 cells, such as gRNA testing for gene knockout.

Our use of FLEx-SaCas9 CFS414 cells in tandem with gRNA-Cre constructs to disrupt endogenous genes is the first demonstration that we are aware of for a CRISPR-Cas9 system in any *Neoaves* species. While CRISPR-Cas9 systems have been used along the phylogenetic tree surrounding the avian clade (Auer and Bene, 2014; Cheng et al., 2018; Doudna and Charpentier, 2014), our test case of disruption of GFP and endogenous genomic DNA sequences substantiate the relative efficacy of Cas9-mediated gene knockout in zebra finch cells. Efficient gRNAs were also found for cutting DNA in multiple genes, expressed at both high and low levels and located on different chromosomes of CFS414 cells. Preliminary validations of cutting in cell lines can provide confidence in investing substantial time and resources to target these genes *in vivo*. The demonstration of SaCas9-mediated genome editing in zebra finches is especially promising for future studies, as SaCas9 may be packaged alongside a gRNA cassette in adeno-associated viruses (AAVs) for local *in vivo* manipulation (Heston and White, 2017; Ran et al., 2015). Other potential applications of CFS414 cells remain to be explored.

Finally, this study highlights the possibility for cell line generation in other non-poultry birds. While there has been evidence of insufficient immortalization from the SV40 large T antigen in other *Neoaves* species (Katayama et al., 2019), our work demonstrates the strength of SV40Tt immortalization in deriving stable zebra finch cell lines. Future work may determine whether this immortalization strategy works in other tissue types and bird species, both within the passerine family and broader avian phylogeny. Overall, cell lines like CFS414 are powerful genetic and molecular resources to expand the application of emerging avian model systems.

## ACKNOWLEDGEMENTS

This work was done in part with help and equipment from Rockefeller University’s Bioinformatics, Flow Cytometry, Genomics, and Bio-Imaging Resource Centers and the Vertebrate Genome Laboratory. The authors would like to thank Samara Brown, Anna Keyte, and Greg Gedman for helpful discussion and manuscript feedback. Special thanks to Jennifer Balacco for help with amplicon library generation, and to Wei Wang for scRNAseq pre-processing. This work was funded through the Howard Hughes Medical Institute and Rockefeller University start-up funding. The pTK608 plasmid was a gift from Tal Kafri (University of North Carolina, Chapel-Hill). The w612-1 plasmid was a gift from Eric Campeau. The pMD2.G and PsPax2 plasmids were gifts from Didier Trono (École Polytechnique Fédérale de Lausanne). The FUGW plasmid was a gift from David Baltimore (California Institute of Technology). The pCMV-PuroR-GFP plasmid was a gift from Jae Yong Han (Seoul National University). The PBCAG-GFP plasmid was a gift from Joseph Loturco (University of Connecticut). The pX601 plasmid was a gift from Feng Zhang (Massachusetts Institute of Technology). The pminiTol2 and pCMV-Tol2 plasmids were gifts from Stephen Ekker (Mayo Clinic).

## AUTHOR CONTRIBUTIONS

M.T.B., and E.D.J conceived the project. M.T.B., P.C., J.M., B.H., generated the sequence libraries. H.U.T. provided equipment, reagents, and expertise. M.T.B. performed experiments. M.T.B. and O.F. analyzed the results. M.T.B. generated the figures. M.T.B., O.F. and E.D.J. wrote the manuscript.

## DECLARATION OF INTERESTS

The authors declare no competing interests.

## DATA AVAILABILITY

## Materials Availability

The CFS414 cell line and select clones will be made available through The Rockefeller University Antibody and Bioresource Resource Center (https://www.rockefeller.edu/monoclonal/cell_line_distribution/), and other clones available upon request. Plasmids generated are available through Addgene or otherwise by request to the corresponding authors.

## Data and Code Availability

Genome assembly data will be submitted to public SRA and NCBI databases. scRNAseq data will be submitted to GEO, and code for Seurat processing and figure generation may be found on Github (http://github.com/Neurogenetics-Jarvis). Accession numbers for public data will be provided upon publication. Requests for datasets generated should be directed toward the corresponding authors.

## METHODS

### Zebra Finch Care and Use

Animals were cared for in accordance with the standards set by the American Association of Laboratory Animal Care and Rockefeller University’s Animal Use and Care Committee. Eggs were collected from single-pair zebra finch nests, with parental coat-color noted, then incubated at 38°C and 50-60% humidity in Showa Furanki incubators (Model #P-018A). For *in* situ hybridization, brain tissue was collected from adult zebra finch males (≥90 days old) as described previously (Biegler et al., 2021).

### Cell collection and culture

Hamburger-Hamilton stage 28 (embryonic day 6) zebra finches (Murray et al., 2013) were removed from eggs and checked for gross anatomical abnormalities. The limbs, head, spine and organs were removed from the embryo, pipetted vigorously in 0.25% Trypsin-EDTA (Gibco, Cat. #25200114) and left to incubate at 37°C for five minutes. The tubes were centrifuged at 200xg for three minutes and resuspended in complete media (DMEM high-glucose, no glutamine (Cat. #11960044) with 10% (v/v) fetal bovine serum (Cat. #21640079), 1x GlutaMAX (Cat. #35050061), 1x Antibiotic-Antimycotic (Cat. #15240062), 1x NEAA (Cat. #1140050), and 1x Sodium pyruvate (Cat. #11360070), each purchased from Gibco™). Undissociated particulates were allowed to sink to the bottom of the tube and the supernatant was plated onto 10cm dishes. Cells were incubated in air-jacketed incubators set to 37°C and 5% CO_2_. All phase-contrast and fluorescent images were taken on a Leica DM 1L inverted microscope unless otherwise specified.

To passage both primary fibroblasts and the CFS414 cells, media was removed at 100% confluence, the monolayer washed with 1xHBSS (Gibco, Cat. #14170112) and incubated in 0.25% Trypsin for 3-5 minutes. Trypsin was neutralized 3:1 with complete media, and the cells were centrifuged at 200xg for five minutes, after which the supernatant was aspirated and the cell pellet resuspended in complete media. Primary fibroblasts were passaged at a split of 1:2 to 1:4 and CFS414 cells were generally passaged at a 1:10 to 1:20 split every 4 to 6 days.

### Cryopreservation

Cells were cryopreserved by mixing 100µL dimethyl sulfoxide (DMSO, Sigma Cat. #D2650) to 900µL complete media containing 2-4 million cells into a cryovial and freezing in a Styrofoam container at -80°C for 24-48 hours, then stored in liquid nitrogen for later use. To thaw, cells were placed in a 37°C water bath until no ice remained in the vial and quickly diluted with 10mL complete media, centrifuged at 200xg for 3 minutes, and then plated onto a 10cm dish with 10mL complete media.

### Growth curves

For growth curve calculation, cells were seeded between 657 cells/cm^2^ (∼1:256 dilution) and 168,421 cells/cm^2^ (∼1:2 dilution) in complete medium in 6 duplicate wells of a 24-well plate (1.9 cm^2^ plating surface). Cell number measurements were performed every 24 hours for at least 6 days after seeding. For cell counting, cells were trypsinized and resuspended in a total volume of 200-1000*µ*L complete media, then counted on a Countess II FL (Applied Biosystems). The log-phase doubling time, *g*, between days was calculated as:

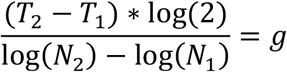

where *T*_2_ and *T*_1_ are the time at collection and the previous collection, and *N*_2_ and *N*_1_ are the averaged cell counts at collection and the previous collection time. Log phase was roughly defined as the 3 steepest consecutive slopes in the growth curve.

### Generation of lentivirus constructs and cell transduction

Plasmids used for lentiviral production may be found in Table S8 alongside their respective sources and RRIDs. Confluent HEK293T (RRID:CVCL_0063) cells were split 1:2 into T75 flasks and were immediately transfected with molar equivalents of pMD2.G, PsPax2 and either pTK608, w612-1, or pMB204 using Lipofectamine 2000 (Invitrogen, Cat. #11668027). After three days, media was collected and passed through a 0.22µm PVDE filter. For pTK608 and w612-1 viral particles, Lenti-X Concentrator (Takara, Cat. #631231) was used following the manufacturer’s protocol and resuspended in HBSS. pMB204 viral particles were spun down at 26,000 rpm for 2 hours and resuspended in HBSS. Viral batches were titrated by serial dilution in HEK293T cultures, calculating approximately 1.0E7 IFU/mL for pTK608, and 1.0E9 IFU/mL for pMB204, while the fluorophore-free w612-1 was assumed to be roughly equivalent to the batch-paired pTK608 titer.

Primary FEFs or CFS414 cells in 24-well plates were transduced with 10-30µL of virus in 300µL of complete media for 6 hours, and then an additional 200µL added to the wells 6 hours later. Media was changed after 48-72 hours.

### *PiggyBac* construct generation and transfection

Information on plasmids used for cloning is found in Table S8 alongside their respective sources and RRIDs. The pMB950 and pMB1052 *piggyBac* FLEx templates were constructed by inserting synthesized gBlocks (Integrated DNA Technologies) into the PBCAG-eGFP backbone digested with *BspDI, SpeI, EcoRI, AgeI*, and/or *NotI* restriction enzymes (purchased from New England Biolabs) using NEBuilder Hi-Fi Assembly (New England Biolabs, Cat. #E2621). The pMB1052 construct was subsequently cut using *BspDI* and *EcoRV* (New England Biolabs), and then cloned into a PCR-amplified pminiTol2 backbone using NEBuilder Hi-Fi Assembly. For pMB952, *S*.*aureus* Cas9 was amplified with homologous adaptors from pX601 by PCR for ligation into the pMB950 plasmid digested with *NheI* and *XhoI* (New England Biolabs).

CFS414 cells were transfected with Lipofectamine 3000 (Invitrogen, Cat. #L3000015) in 12-well or 6-well plates, using manufacturer’s instructions. Using vectors with antibiotic selection, cells were washed after 48-72 hours after transfection with complete media containing 0.5µg/mL Puromycin (Gibco, Cat. #A1113802) or 1.5mg/mL Geneticin (Gibco, Cat. #10131035), with antibiotic media replaced every three days until the desired selection level was reached. *PiggyBac* transposon vectors were co-transfected with the pCyL43 *piggyBac* transposase vector (Li et al., 2013). Tol2 transposon vectors were co-transfected with pCMV-Tol2.

### Genomic loci amplification

Genomic DNA was extracted from cells by lysis in WTL buffer (25mM Tris-HCl pH 8.0, 10mM EDTA pH 8.0, 1% SDS), treatment with 15µg RNAse A, protein precipitation in 7.5M ammonium acetate, followed by isopropanol and ethanol precipitation prior to DNA resuspension in Ultrapure distilled water (Invitrogen, Cat #10977015). Genomic loci were then amplified using primers targeting genes of interest (see Table S9 for a list of primers) using Q5 Hot Start High-fidelity 2X Master Mix (NEB, Cat. #M0494). One exception was the *CHD* sextyping amplification protocol, which followed a Taq polymerase protocol from a previous study (Griffiths et al., 1998).

### Karyotyping

CFS414 cells were seeded at 1:4 dilution in T-25 flasks and shipped overnight to KaryoLogic, Inc. (Durham, NC), where Giemsa-banded karyotyping was performed as previously described (Rosselló et al., 2013).

### Pacbio gDNA sequencing, assembly, and structural variant analysis

Roughly 4 million CFS414 cells were trypsinized, washed in 1xDPBS, and centrifuged before storing the pellet at -80°C. Ultra-high molecular weight (uHMW, ∼50-300 kb) DNA was extracted using the Nanobind CBB Big DNA Kit (Cells, Bacteria, Blood) (Circulomics, Cat. #NB-900-001-01). uHMW DNA quality was assessed by a Pulsed Field Gel assay and quantified with a Qubit 2 Fluorometer, and 5µg of uHMW DNA was sheared using a 26G blunt end needle (Pacbio protocol PN 101-181-000, Version 05). A large-insert (23 kb average) Pacbio library was prepared using the Express Template Prep Kit v1.0 (Pacific Biosciences, Cat. #101-357-000) following the manufacturer’s instructions. Library size selection using the Sage Science BluePippin Size-Selection System was made for inserts larger than 20kb. This library was sequenced with a Pacific Biosciences 8M (Cat. #101-820-200) SMRT cell on the Sequel II instrument with the Sequencing Kit 2.0 (Cat. #101-820-200) and Binding Kit 2.0 (Cat #101-842-900), recording a 15-hour movie of sequence reads. Raw Pacbio subreads were mapped to the zebra finch reference genome bTaegut1.v2 (with W chromosome from bTaegut2.pat added) using minimap2 (Li, 2018). Base coverage was calculated using SAMtools 1.9 (Li et al., 2009) and BEDtools genomecov v2.27.1 (Quinlan and Hall, 2010).

Pacbio subreads were filtered using BLAST (Altschul et al., 1990). Subreads that have a significant overlap with the w612-1 proviral sequence were selected and mapped to the reference genome using minimap2 and bam files were sorted and indexed using SAMtools. Alignment was visualized in IGV (Robinson et al., 2011) and plotted using R custom scripts.

A total of 139,28,268,101 bp of sequence was generated with a subread N50 of 33,751bp. The genome was assembled using Falcon-unzip 2.2.4 (Chin et al., 2016), with primary contigs filtering for artifacts (haplotypic duplications) using Purge-dups (Guan et al., 2020). The final primary assembly length was 1,047,588,919bp, composed of 356 contigs (N50 contig= 12,971,239bp).

### Bionano structural variant analysis

uHMW DNA was labeled for Bionano Genomics optical mapping using the Bionano Prep Direct Label and Stain (DLS) Protocol (Cat. #30206E). Labeled DNA was loaded and run on one Saphyr instrument chip flowcell for optical imaging. 420.67 Gb (read length ≥150 kb) was generated with read length N50 = 343.5 kb and average Label density of 13.92 labels per 100kb. A consensus genome map was constructed and aligned to the zebra finch reference genome using Solve 3.6.1_11162020. The Variant Annotation Pipeline was used to identify signatures of structural variants.

### Single-cell RNA sequencing

CFS414 cells at passage 24 and passage 50 were trypsinized and counted on a Countess II FL (Life Technologies), and a single cell suspension was diluted to around 1,000 cells/µL by centrifugation at 200xg and resuspension in complete media. Approximately 5,000 cells were loaded into a Chromium Chip (10x Genomics). cDNA synthesis and library preparation were performed using the Chromium Single Cell 3’ Reagent Kits (version 3, 10x Genomics) according to manufacturer’s instructions. Sequencing was performed by Novogene Co., Ltd. on an Illumina HiSeq 4000.

Raw sequencing data was aligned to a VGP male zebra finch genome assembly (bTaegut1.v2), with the addition of the W chromosome from bTaegut2.pat.W, using the CellRanger analysis pipeline (10x Genomics, version 3). The 3’ and 5’ UTRs of annotated genes were extended using previously collected bulk RNAseq data to enhance read alignment. Analysis of the sequence data was performed using the Seurat tools workflow (version 3.2.0) (Butler et al., 2018) in RStudio (version 4.0.2). Briefly, datasets were trimmed, normalized and scored for cell cycling to regress zebra finch genes and Refseq IDs associated with G1, G2M and S phases, as well as mitochondrial genes. These datasets were then scaled by SC-Transform (Hafemeister and Satija, 2019) using Seurat commands. The two datasets were integrated using anchor-based nearest neighbor functions using default settings, and 50 PCAs were used to calculated. Given the sequence depth and strong homogeneity of the samples, a selection of 13 PCA dimensions for UMAP generation and Nearest Neighbor clustering was chosen to avoid over-segregation by residual cell cycling and ribosomal protein variability. A clustering resolution, visualized using the clustree tool (version 0.4.3, https://CRAN.R-project.org/package=clustree), was similarly chosen at 0.3 to avoid oversplitting by these gene categories. Hierarchical clustering was estimated using the FindClusterTree function (dimensions = 10) in Seurat. Average gene expression and expressing population percentage was extracted from the metadata of the unscaled DotPlot function. LogFC, standardized variability, and passage marker p-values were appended to the table using the metadata from the VariableGenesPlot and FindMarkers functions (set.ident = “passage”). GO terms for the gene markers were found using the GOrilla webtool (Eden et al., 2009).

### Fluorescence-activated cell sorting and clonal propagation

Cells were dissociated with 0.25% Trypsin, pelleted at 200xg for 3 minutes, and resuspended in cell sorting buffer (1x DPBS (Ca/Mg++, pH 7.0-7.4), 10mM HEPES (Gibco, Cat. #15630080), pH 7.0, 0.1% Bovine Serum Albumin, (Invitrogen, Cat. #AM2616), 5mM EDTA, and 2-8 ng/mL DAPI (4′,6-diamidino-2-phenylindole, used as a vital stain)), passed through a 40µm filter and diluted to 1.0-3.0E6 cells/mL. Cells were taken to the Rockefeller Flow Cytometry Resource Center and loaded into a FACS Aria II (BD Biosciences). Live (DAPI-free) cells were sorted by desired fluorescent intensity. Samples were either sorted in bulk (purified) or single (clonal) cells, index-sorted into a 96-well plate containing 100µL conditioned, filtered complete media from CFS414 cells.

For purified samples, the sorted cells were washed twice with complete media with 5x antibiotic-antimycotic and plated onto a 10cm dish. For clonal samples, plates were incubated at 37°C, 5% CO_2_. An additional aliquot of complete media (50µL) was added 72 and 168 hours after sorting. After 11 days, colony formation was assessed on an inverted microscope. Suitable clones were harvested for expansion after reaching an adequate colony size for passaging (around 1/4^th^ of the well surface).

### Single-label *in situ* hybridization

Digoxigenin-(DIG) conjugated probes for *ZEB2* (Genbank: DV946921) and *PVALB* (Genbank: DQ215755) were made and applied onto male zebra finch brain sections from previous studies, as detailed previously (Biegler et al., 2021). For the *HTR1A* DIG-labeled probe, the sequence was amplified from zebra finch cDNA using primers containing the T7 and T3 promoters (Table S9), and the probe was generated and used in the same way as the others.

### GFP-SAP30L cytoplasmic shuttling assay

pMB204-transduced cells were plated onto coverslips and challenged with 0.0 mM or 1.0 mM H_2_O_2_ in fresh culture media for 15 minutes. Conditioned media was then replaced and the cells incubated at 37°C. After 4 hours, cells were washed with 1xPBS and fixed for 15 minutes in in 4% PFA in 1xPBS. DAPI nuclear counterstain was conducted and the slides coverslipped with ProLong Diamond Antifade mounting medium.

### Zebra finch primary neuron culture

For the culture of zebra finch neurons, an adapted protocol of mouse cortical neuron culture was used (McDowell et al., 2010). Briefly, Hamburger-Hamilton stage 45 embryos or day 0 hatchlings were euthanized, followed by brain dissection and the removal of meninges. Hemispheres were approximately resected to remove the basal ganglia (Chen et al., 2013) and the cortical regions were placed in 200 arbitrary units/mL papain (Worthington, Cat. #3126) for 5-7 minutes and neutralized with trypsin neutralization solution (Sigma, Cat. #T9253). Neurons were then counted and plated onto Laminin-treated coverslips (Neuvitro, Cat. #GG-12-Laminin) at a density of 50,000 cells/cm^2^ in cortical plating media (Basal Medium Eagle (Sigma B1522), 0.6% Glucose, 10% (v/v) fetal bovine serum, 1x GlutaMAX, 1x Antibiotic-antimycotic). One day after plating, media was removed and replaced with serum-free neurobasal media (Gibco, Cat. #21103049) with 1x B27 supplement (Gibco, Cat. #17504044), 1x GlutaMAX, and 1x Antibiotic-antimycotic.

Neurons were transfected at 6 days *in vitro* (DIV6) with pMB204 plasmid using a mixture of Lipofectamine 3000 with the Combi-Mag reagent (OZ Biosciences, Cat. #KC30200) according to manufacturer’s instructions. This provided strong and consistent expression in the transfected, post-mitotic cells. At DIV8, cells were treated with cyanquixaline (CNQX, Sigma, Cat. #C127), Bicuculline (Sigma, Cat. #14340), tetrodotoxin (TTX, Abcam, Cat. #ab120054), and DL-2-amino-5-phosphonopentanoate (APV, Sigma, Cat. #A5282) (Table S7). After 16 hours, neurons were challenged with fresh media, 1mM H_2_O_2_ for 15 minutes, or 50µM NMDA for 1 hour, then allowed to recover for four hours in their corresponding condition’s media before being fixed with 4% PFA in 1xPBS for 15 minutes.

### Immunocytochemistry

Neurons on coverslips were fixed with 4% ice-cold PFA, washed with 1xPBS, and permeabilized with 0.5% Triton-X in 1xPBS. Prepared slides were then blocked in 10% Blocking One (Nacalai, Cat. #0395395) with 0.5% Triton-X and 1xPBS for 30 minutes, then incubated with 1:1000 primary antibody (Chicken anti-GFP (Sigma, Cat. #AB16901, RRID:AB_11212200) and Mouse anti-MAP2 (Sigma, Cat. #MAB3418, RRID:AB_94856) overnight at 4°C. Sections were then washed 3 times with 1xPBS with 1% Tween-20 (PBS-T) for 15 minutes and then transferred to host-specific 1:200 fluorophore-conjugated secondary antibody (Goat anti-Chicken 488 (ThermoFisher, Cat. #A11039, RRID:AB_142924) and Goat anti-Mouse (ThermoFisher, Cat. #A21125, RRID:AB_141593)) for one hour at RT. Cells were then washed again with 1xPBS-T and DAPI nuclear counterstain was applied before mounting media and coverslip were placed on slides for imaging.

### CRISPR-gRNA cutting assays

The LacZ and GFPg1 gRNAs were selected for minimal zebra finch genome sites for potential off-target cutting using GT-Scan (O’Brien and Bailey, 2014). For gRNA sequences, see Figure S7. Oligonucleotides (Integrated DNA Technologies) of the gRNA sequence were then annealed and ligated into the pMB1052 vector, using a protocol for the AAV construct, pX601 (https://www.addgene.org/crispr/zhang/). Cells were lipofected with pMB952, pMB1052, and pCyL43 and purified with Geneticin 48 hours later (Gibco, Cat. #10131027). Prior to imaging, cells were washed with complete media containing 100ng/mL Hoescht33342 (Invitrogen, Cat. #H3570).

Endogenous genes were targeted using gRNAs designed by GT-Scan or CRISPOR (Concordet and Haeussler, 2018). CFS414 clone 1G10 cells with the transposon-integrated pMB952 construct were transfected with pMB1052 constructs containing gRNAs targeting endogenous zebra finch genes (for sequences, see Figure 7C) using Lipofectamine 2000. Cells were selected with 1.5mg/mL Geneticin (Gibco, Cat. #10131035) and further purified by bulk and monoclonal FACS, as outlined above. We noted GFP and mCherry signal was variable, potentially due to genomic interference or loss of *piggyBac* integration. Genomic DNA was extracted, and the genomic loci surrounding the gRNA or predicted off-target sites was amplified by PCR (see Table S9 for primers). Illumina N7xx and N501 adaptors were added by further PCR amplification (Joung et al., 2017). Primers were removed using AMPure XP beads (Beckman Coulter, Cat. #A63880), and the amplicon quality was assessed on an Agilent 2100 BioAnalyzer before samples were pooled into a 10µM DNA library. DNA libraries were sent to the Rockefeller University Genomics Resource Center to run on a MiSeq benchtop sequencer (Illumina) according to manufacturer’s instructions. From these amplicon-derived sequences, indel formation was assessed and visualized using CRISPResso2 (Pinello et al., 2016).

### Quantification and statistics

All genomic, transcriptomic, and amplicon dataset quantification and statistical analysis was performed as described above.

For the CFS414 cytoplasmic shuttling assay, samples were imaged on a Zeiss LSM 780 laser scanning confocal microscope. To determine cell localization, non-transduced cells (<1% total population) were identified and used to normalize for background signal in the image (n = 4 images per condition). Images were blinded for analysis, and DAPI signal was subtracted from GFP signal in transfected cells, and cytoplasmic GFP signal in the remaining cells was counted. Significance was determined using a Student’s two-tailed, equal variance t-test.

To determine cell localization of GFP in primary neurons, several images were collected per condition using an Olympus BX61 upright microscope. Images (n= 6-8 images per condition) were blinded for analysis, and GFP signal co-localized with blue DAPI signal and surrounded by MAP2 cytoskeletal marker was identified as neuronal expression. An average of 9.7 transfected neurons were identified in each image. GFP signal that extended beyond the DAPI-stained nucleus and into the MAP2-stained processes was considered cytoplasmic localization. A percentage of cytoplasmic localization was calculated per image and averaged for each condition. Significance was calculated with a one-factor ANOVA with a post-hoc Tukey-Kramer test.

For the CFS414-GFP clone 2D4 gRNA targeting experiments, cell nuclei were counted in FIJI (n = 4 images per condition), then co-localized with GFP signal to determine percentage of total cells. GFP signal loss was visibly apparent without quantification between conditions, so this analysis was unblinded. Significance was determined by a Student’s two-tailed, equal variance t-test.

## SUPPLEMENTAL FIGURES

**Figure S1.**
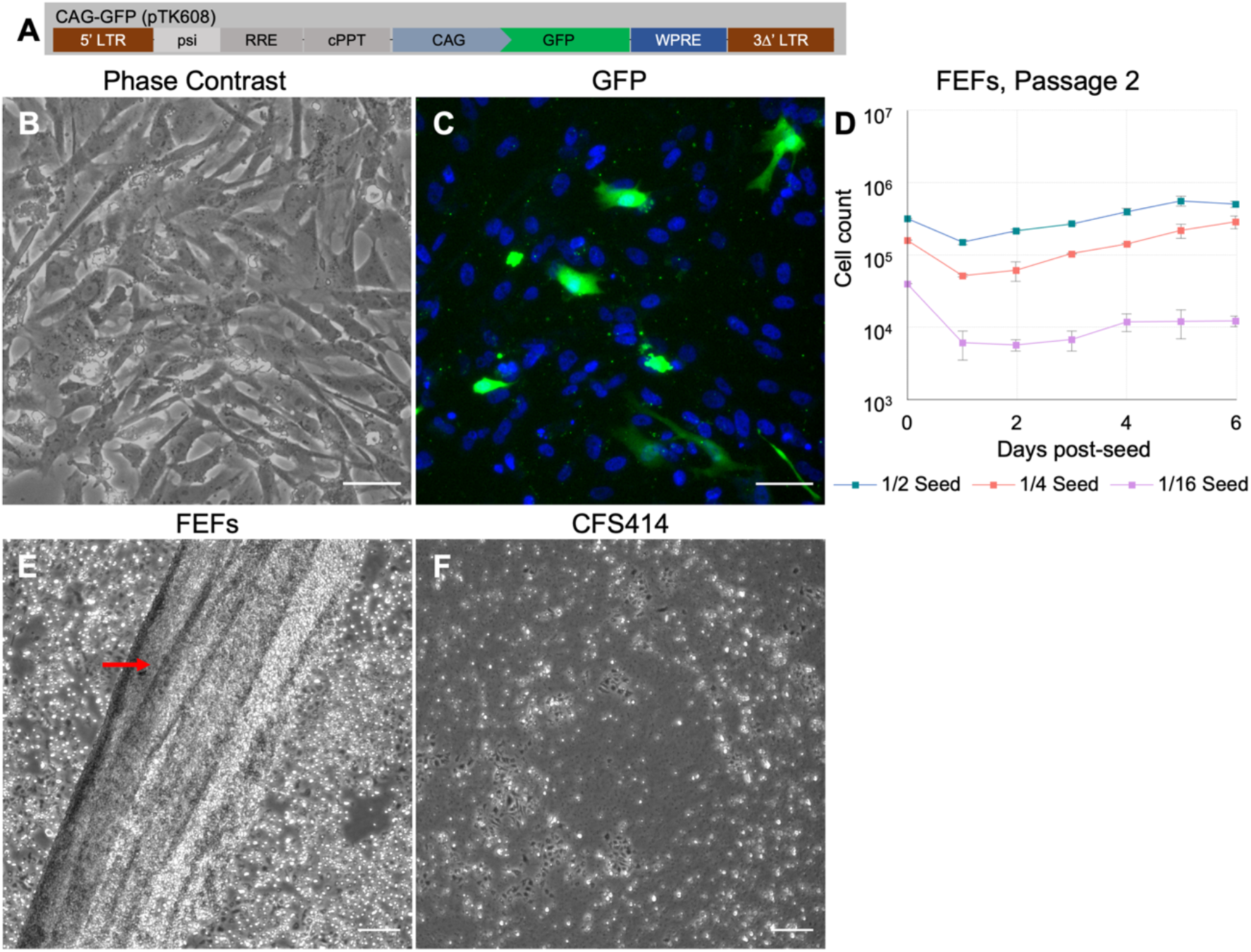
Zebra finch embryonic fibroblast characterization. (A) Diagram of CAG-GFP control lentiviral construct. (B-C) Passage 2 Finch embryonic fibroblasts (FEFs) 72 hours following transduction with CAG-GFP (pTK608) control lentivirus imaged by (B) phase-contrast and (C) fluorescence to show sparse GFP transduction with Hoescht33342 nuclear counterstain. Scale bars = 50µm. (D) Growth curve of passage 2 FEFs at variable seeding densities, ranging from 1:2 (320,000 cells/well) to 1:16 (40,000 cells/well) dilution. Note the difference in growth between seeding densities compared to CFS414 cells (Figures 1G and 1H). Error bars denote SEM. (E) Exemplary image of collagenated sheet formation (red arrow) by overconfluent FEFs that have been trypsinized. (F) Similarly overconfluent, trypsinized CFS414 cells for comparison. Scale bars (E-F) = 200*µ*m.

**Figure S2.**
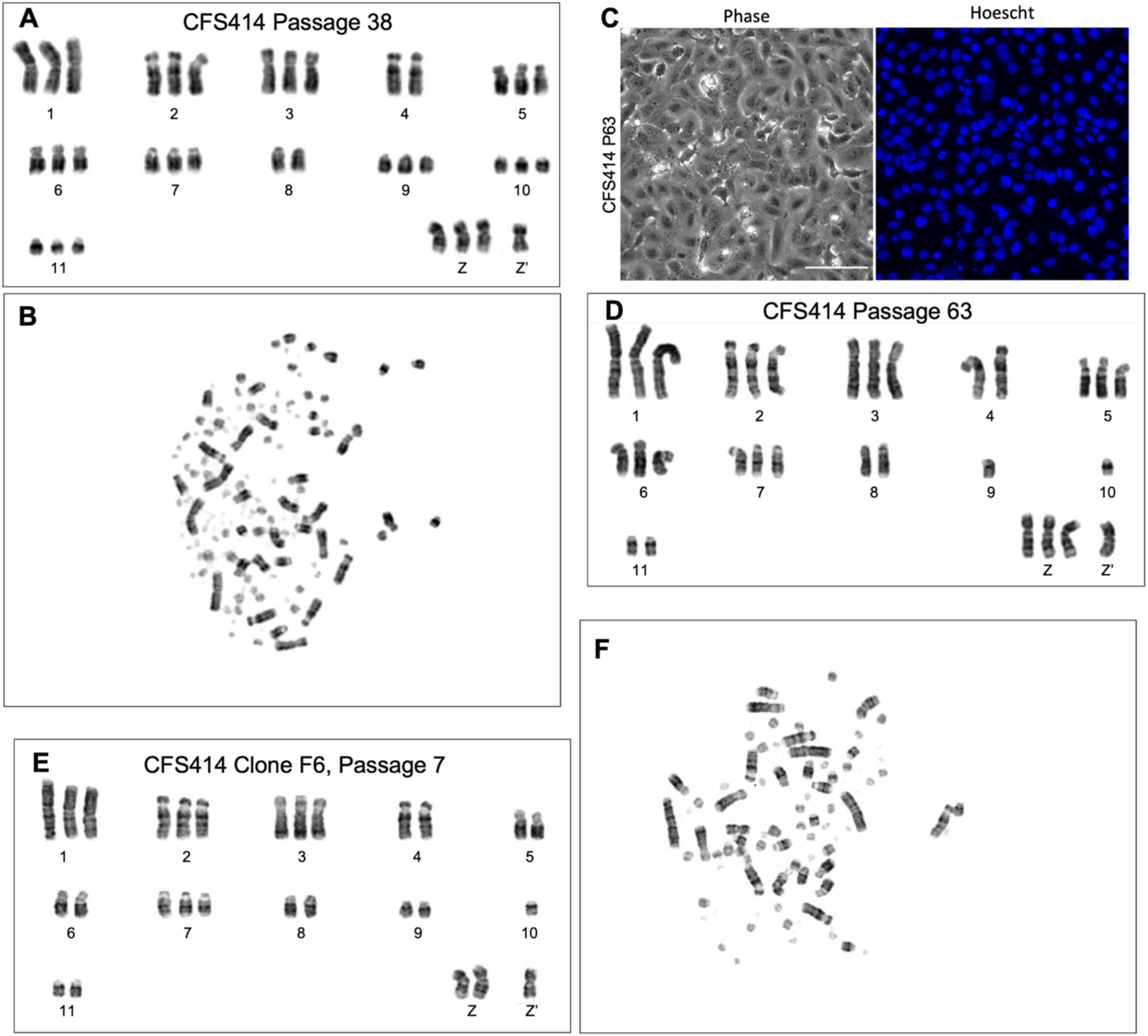
Karyotypes of CFS414 passage and clonal cell populations. (A) Second exemplar karyotype of passage 38 CFS414 cells. (B) Corresponding metaphase spread for (A). Note the presence of additional microchromosomes. (C) Phase contrast and Hoescht33342-stained images of passage 63 CFS414 cells. Scale bar = 50µm. (D) Example karyotype of passage 63 CFS414 cells. (E) Second exemplar karyotype of CFS414 clone F6 cells with (F) Corresponding metaphase spread for (E). Note presence of mutant Z chromosome (Z’) in all karyotypes.

**Figure S3.**
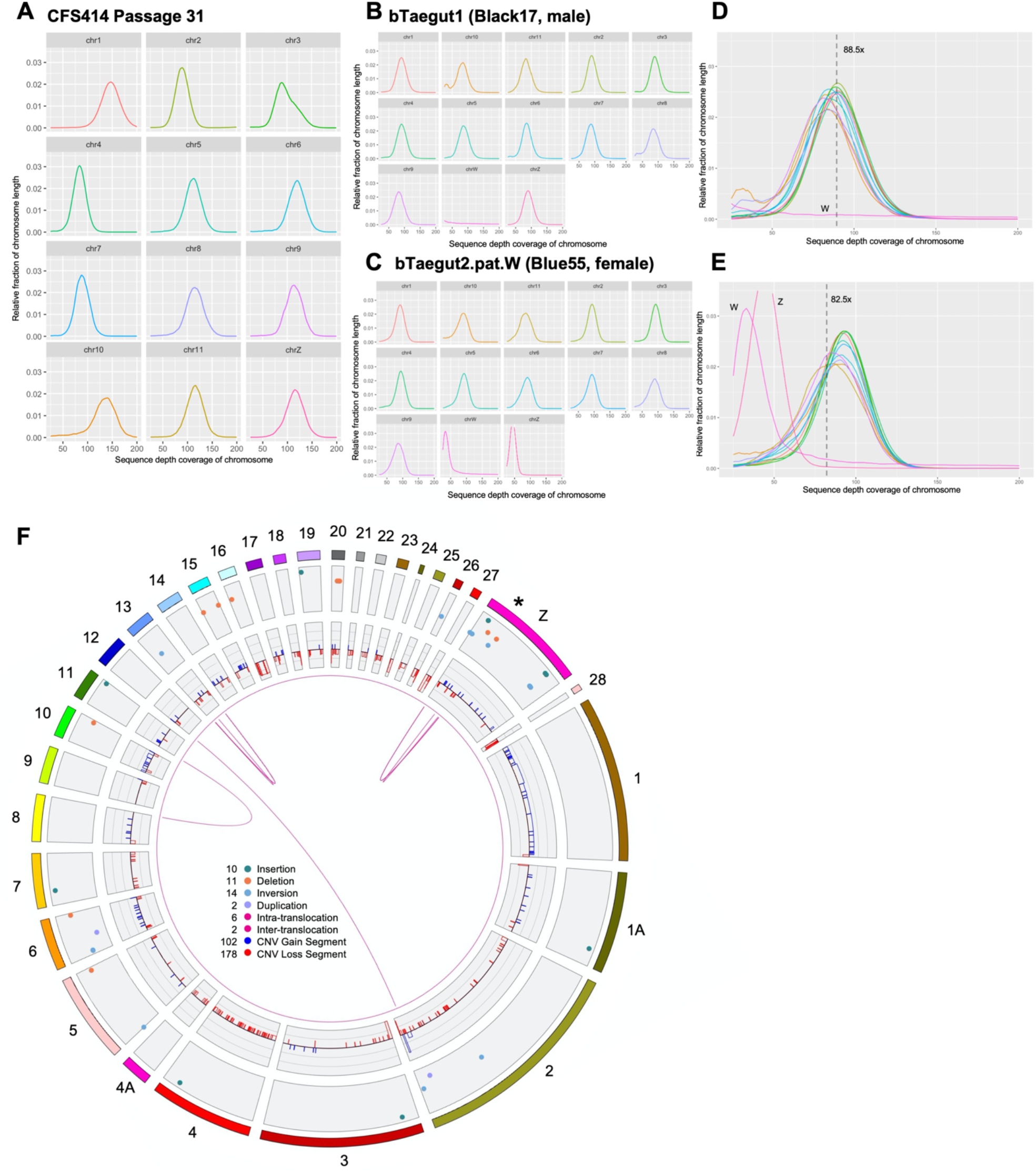
Structural variants within the CFS414 genome assembly. (A) Fraction of coverage for each CFS414 macrochromosome, separated out from Figure 2F. (B-E) Fraction of coverage for each chromosome in (B) bTaegut1 and (C) bTaegut2.pat.W male and female reference genomes, respectively, overlaid in (D) and (E) as single charts. Dotted lines denote the sequence coverage for each assembly. (F) Circos plot of CFS414 genome assembly compared to the bTaegut1 reference genome, showing large (≥150kb) CFS414 structural variants, specified by the color legend in the center. * denotes highly variable region between the CFS414 and bTaegut1 Z chromosome.

**Figure S4.**
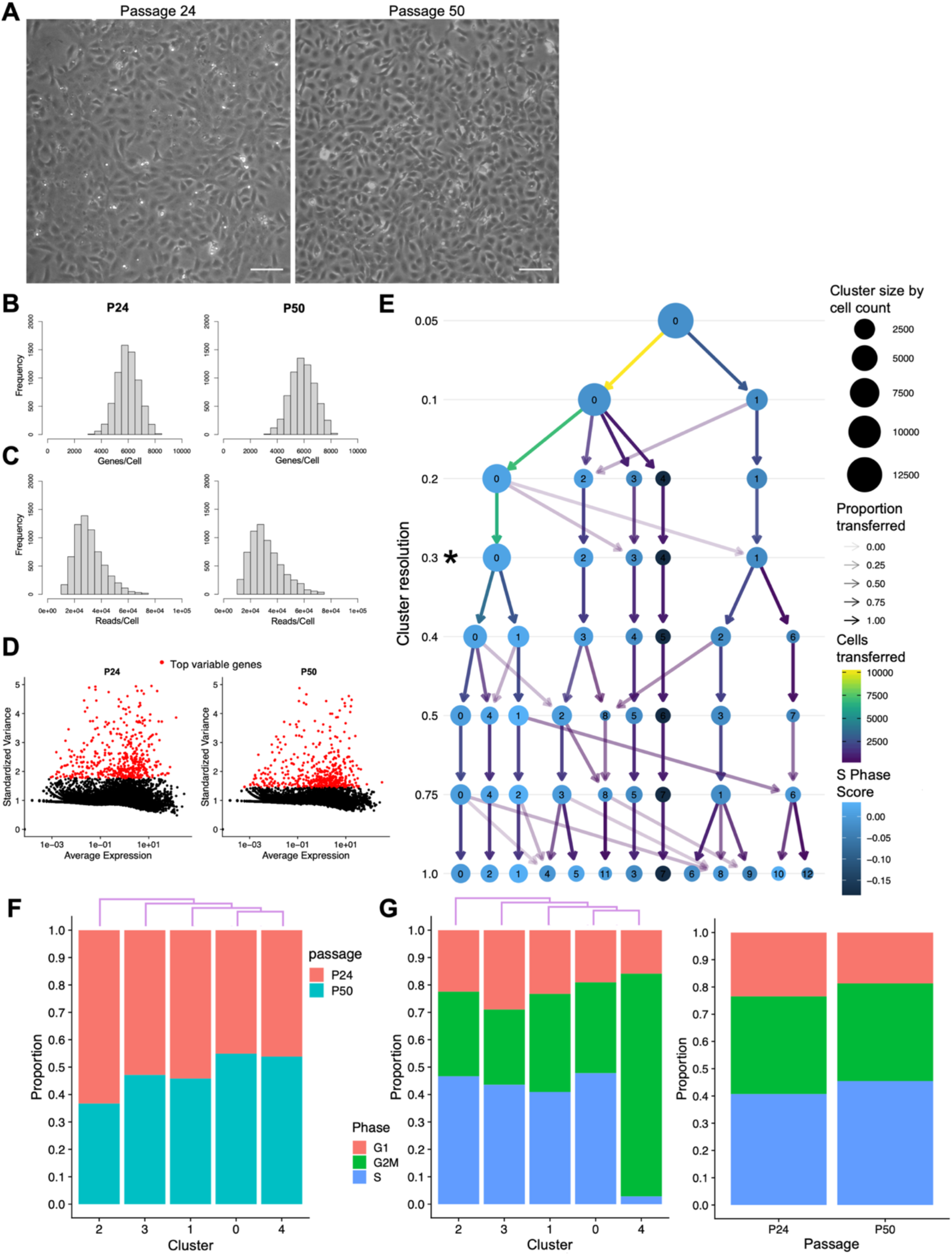
Single-cell RNA sequencing data. (A) Phase contrast images of CFS414 cell samples prior to trypsinization for scRNAseq. Scale bars = 100µm. (B-C) Histograms of (B) Feature counts (Genes/cell) and (C) Raw RNA counts (Reads/cell) for passage 24 (P24, left) and passage 50 (P50, right) CFS414 cell samples. (D) Plot of all annotated genes by standardized variance (y-axis) and average expression (x-axis) for P24 (left) and P50 (right). Subset of Figure 3B (y-axis from 0 to 5) to emphasize difference in top genes by standardized variance (red) between passages. (E) Cluster tree of scRNAseq object for cluster resolution selection, showing cluster formation and stability over resolution values. Cluster nodes are colored by S Phase cell cycle score. * highlights chosen resolution (0.3) for downstream analysis based on cluster stability and cell cycle homogeneity. (F) Proportional bar chart of cluster makeup by passage, with cluster hierarchy defined by differential gene expression for each cluster in purple. (G) Proportional bar chart of cell cycle phase (determined through scRNAseq processing), split by cluster (left) and passage (right). Abbreviations: G1, Growth 1 phase; G2M, Growth 2 and Mitotic phases; S, synthesis phase.

**Figure S5.**
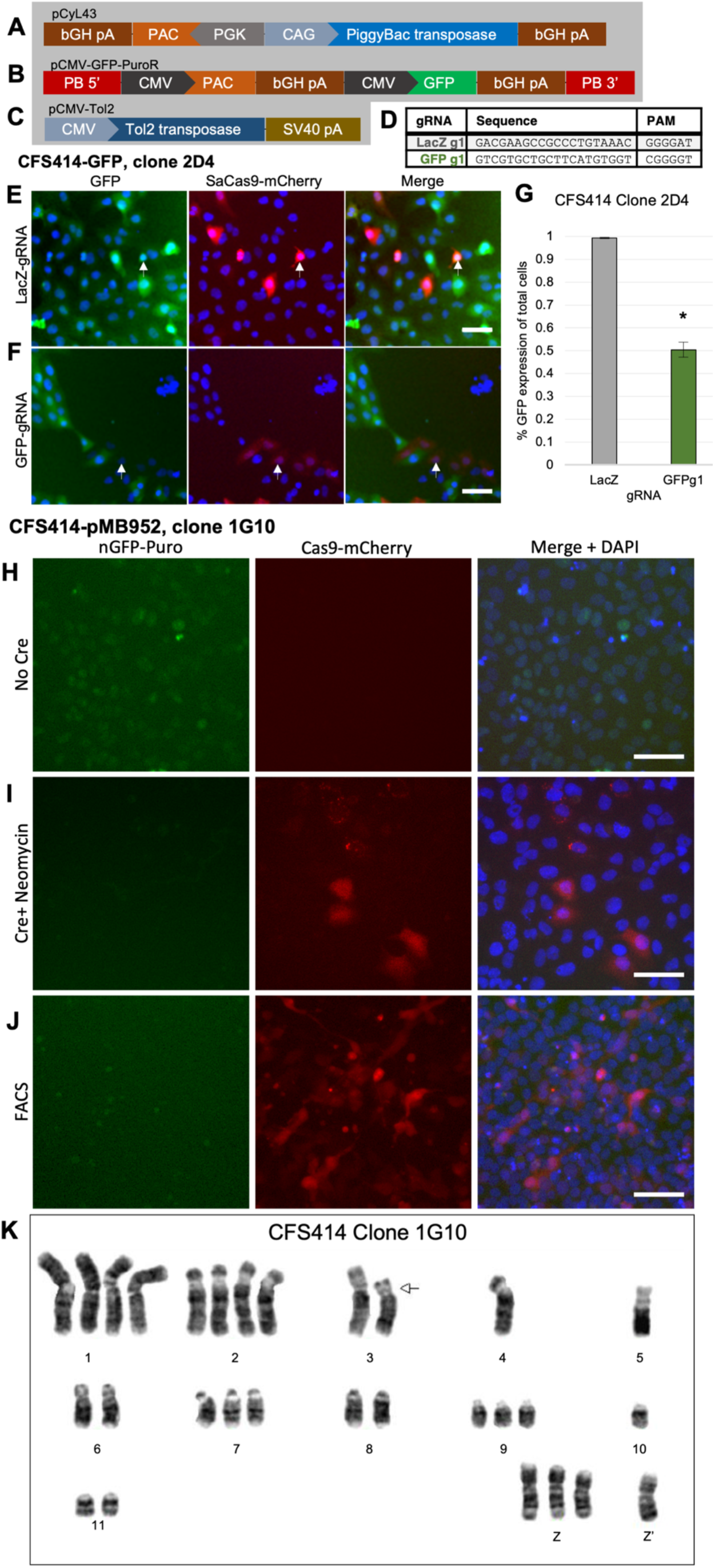
Function of pMB952 and pMB1052 vectors. (A) *PiggyBac* transposase vector pCyL43. (B) *PiggyBac* transposon vector expressing GFP and the Puromycin resistance gene, Puromycin N-acetyltransferase (PAC). (C) *Tol2* transposase vector pCMV-Tol2 for transfection with pMB1052, Figure 6B. (D) gRNAs used to target GFP or LacZ genes with respective protospacer adjacent motif (PAM) listed. (E-G) The monoclonal GFP+ cell line CFS414-GFP clone 2D4, transfected with the *piggyBac* transposase, pMB952, and pMB1051 with (E) LacZ or (F) GFP gRNAs. (G) Percent cells in LacZ and GFPg1 groups expressing GFP post-selection. * denotes significance from a two-tailed t-test, p < 0.05. Error bars denote SEM. (H-J) CFS414 clone 1G10 cells integrated with the pMB952 plasmid. (H) Nuclear green (nGFP) signal from the unrecombined pMB952 plasmid, overlaid with Hoescht33342 nuclear stain (blue). (I) Cre expression through pMB1052 transfection into some cells conditionally excised nGFP signal and expressed SaCas9-P2A-mCherry. Geneticin treated to reduce untransfected cells. (J) FACS-mediated population of 1G10 cells, showing high mCherry populations. (K) Exemplary karyotype of CFS414 clone 1G10 cells at passage 9. Arrow denotes partial deletion in chromosome 3. Scale bars = 50µm.

**Figure S6.**
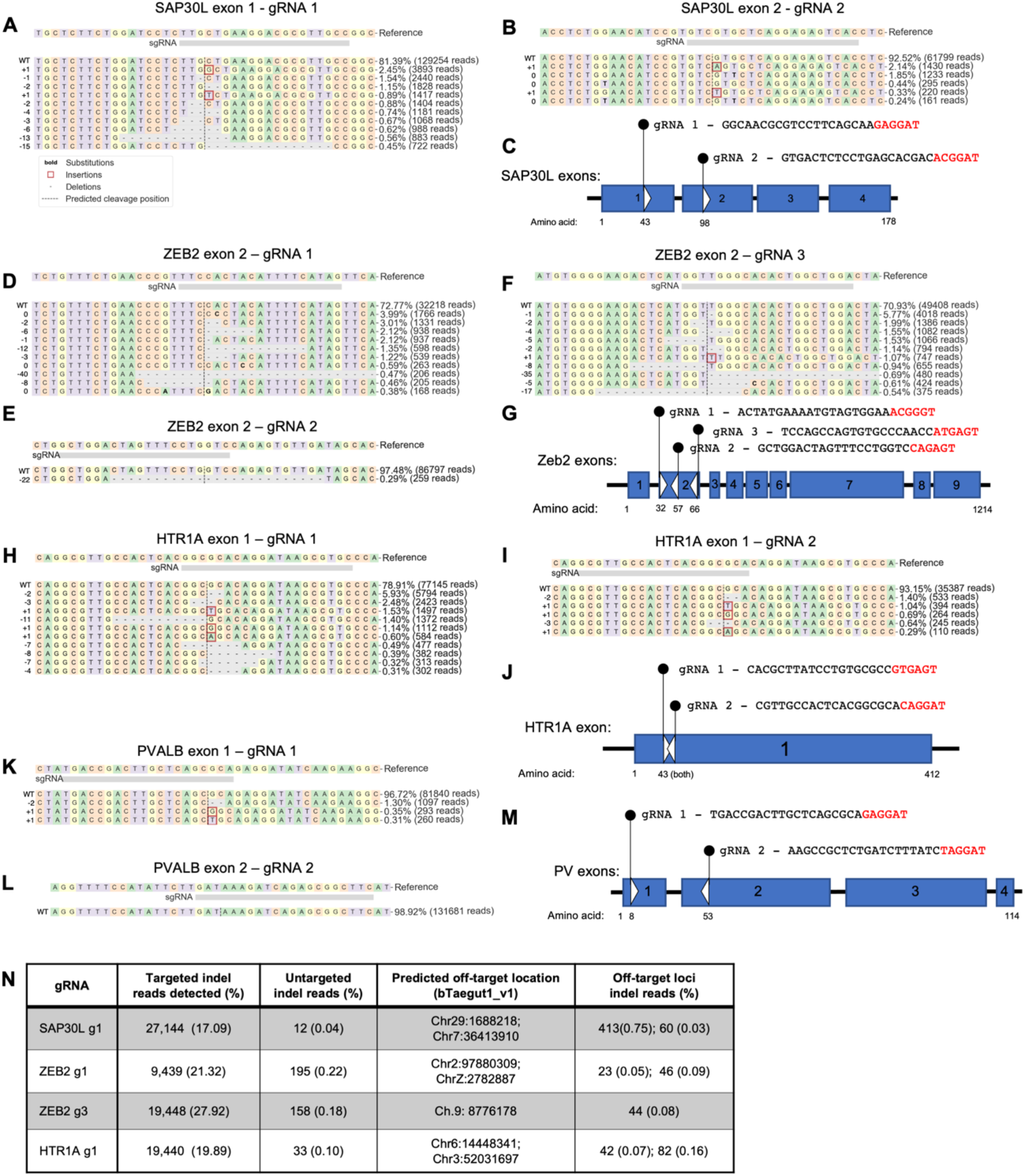
Top CFS414 indel formations from gene targeting gRNAs. (A-B) Sequence reads of *SAP30L* loci following targeting by SAP30L (A) exon 1 gRNA 1 and (B) exon 2 gRNA 2. Legend below (A) denotes indel formation type. (C) Diagram of gRNA target sites on the *SAP30L* gene, showing the exon and predicted amino acid codon sequence targeted. (D-F) Sequence reads of *ZEB2* loci following targeting on exon 2 by (D) gRNA 1, (E) gRNA 2 and (F) gRNA 3. (G) Diagram of gRNA target sites on the *ZEB2* gene, showing the exon and predicted amino acid codon sequence targeted. (H-I) Sequence reads of *HTR1A* locus following targeting by (H) gRNA 1 and (I) gRNA 2. (J) Diagram of gRNA target sites on the *HTR1A* gene, showing the exon and predicted amino acid codons targeted. (K-L) Sequence reads of *HTR1A* locus following targeting by (K) gRNA 1 and (L) gRNA 2. (M) Diagram of gRNA target sites on the *PVALB* gene, showing the exon and predicted amino acid codons targeted. All gRNAs show top 10 indel sequences found in more than 0.3% of reads. (N) Table of indel sequence read counts and their relative percent of total reads for the 4 most fficient gRNAs. Table shows targeted and untargeted CFS414 clone 1G10 samples, and gRNA-targeted 1G10 cells at their respective predicted off-target loci.

## SUPPLEMENTAL TABLE LEGENDS

**Table S1**. CFS414 genome sequencing and assembly statistics **Table S2**. Single-cell RNA sequencing (scRNAseq) statistics **Table S3**. Barcode information for scRNAseq

**Table S4**. CFS414 scRNAseq expression data for all annotated zebra finch genes

**Table S5**. Gene markers for each CFS414 passage 24 scRNAseq cell cluster

**Table S6**. GO Term analysis of human orthologs of CFS414 passage 24 scRNAseq cluster gene markers

**Table S7**. Strategy used for neuronal GFP-TgSAP30L localization experiment

**Table S8**. Plasmids used in this study

**Table S9**. Primers used in this study

## Notes

### Competing Interest Statement

The authors have declared no competing interest.

## References

Ahmadiantehrani, S., and London, S.E. (2017). A reliable and flexible gene manipulation strategy in posthatch zebra finch brain. Scientific Reports 7, 43244–16.

Altschul, S.F., Gish, W., Miller, W., Myers, E.W., and Lipman, D.J. (1990). Basic local alignment search tool. Journal of Molecular Biology 215, 403–410.

Auer, T.O., and Bene, F.D. (2014). CRISPR/Cas9 and TALEN-mediated knock-in approaches in zebrafish. Nature Methods 69, 142–150.

Baba, T.W., Giroir, B.P., and Humphries, E.H. (1985). Cell lines derived from avian lymphomas exhibit two distinct phenotypes. Virology 144, 139–151.

Balakrishnan, C.N., Lin, Y.-C., London, S.E., and Clayton, D.F. (2012). RNA-seq transcriptome analysis of male and female zebra finch cell lines. Genomics 100, 363–369.

Batsché, E., Lipp, M., and Cremisi, C. (1994). Transcriptional repression and activation in the same cell type of the human c-MYC promoter by the retinoblastoma gene protein: antagonisation of both effects by SV40 T antigen. Oncogene 9, 2235–2243.

Biegler, M.T., Cantin, L.J., Scarano, D.L., and Jarvis, E.D. (2021). Controlling for activity‐dependent genes and behavioral states is critical for determining brain relationships within and across species. J Comp Neurol.

Brainard, M.S., and Doupe, A.J. (2013). Translating birdsong: songbirds as a model for basic and applied medical research. Annual Review of Neuroscience 36, 489–517.

Bryson-Richardson, R.J., and Currie, P.D. (2008). The genetics of vertebrate myogenesis. Nat Rev Genet 9, 632–646.

Butler, A., Hoffman, P., Smibert, P., Papalexi, E., and Satija, R. (2018). Integrating single-cell transcriptomic data across different conditions, technologies, and species. Nature Biotechnology 36, 411–420.

Chang, H., and Delany, M.E. (2004). Karyotype stability of the DT40 chicken B cell line: macrochromosome variation and cytogenetic mosaicism. Chromosome Research 12, 299–307.

Chen, C., Winkler, C.M., Pfenning, A.R., and Jarvis, E.D. (2013). Molecular profiling of the developing avian telencephalon: Regional timing and brain subdivision continuities. J Comp Neurol 521, 3666–3701.

Cheng, Y., Lun, M., Liu, Y., Wang, H., Yan, Y., and Sun, J. (2018). CRISPR/Cas9-Mediated Chicken TBK1 Gene Knockout and Its Essential Role in STING-Mediated IFN-β Induction in Chicken Cells. Frontiers in Immunology 9, 3010.

Chin, C.-S., Peluso, P., Sedlazeck, F.J., Nattestad, M., Concepcion, G.T., Clum, A., Dunn, C., O’Malley, R., Figueroa-Balderas, R., Morales-Cruz, A., et al. (2016). Phased diploid genome assembly with single-molecule real-time sequencing. Nature Methods 13, 1050–1054.

Choe, H.N., and Jarvis, E.D. (2021). The role of sex chromosomes and sex hormones in vocal learning systems. Horm Behav 132, 104978.

Concordet, J.-P., and Haeussler, M. (2018). CRISPOR: intuitive guide selection for CRISPR/Cas9 genome editing experiments and screens. Nucleic Acids Res 46, gky354..

Condro, M.C., and White, S.A. (2014). Distribution of language‐related Cntnap2 protein in neural circuits critical for vocal learning. J Comp Neurol 522, 169–185.

Ding, L., and Perkel, D.J. (2004). Long-Term Potentiation in an Avian Basal Ganglia Nucleus Essential for Vocal Learning. J Neurosci 24, 488–494.

Doudna, J.A., and Charpentier, E. (2014). The new frontier of genome engineering with CRISPR-Cas9. Science 346, 1258096–1258096.

Drobik-Czwarno, W., Wolc, A., Fulton, J.E., Jankowski, T., Arango, J., O’Sullivan, N.P., and Dekkers, J.C.M. (2018). Genetic basis of resistance to avian influenza in different commercial varieties of layer chickens. Poultry Sci 97, 3421–3428.

Eden, E., Navon, R., Steinfeld, I., Lipson, D., and Yakhini, Z. (2009). GOrilla: a tool for discovery and visualization of enriched GO terms in ranked gene lists. Bmc Bioinformatics 10, 48.

Ericson, P.G.P., Irestedt, M., and Johansson, U.S. (2003). Evolution, biogeography, and patterns of diversification in passerine birds. J Avian Biol 34, 3–15.

Fitzsimmons, R.E.B., Mazurek, M.S., Soos, A., and Simmons, C.A. (2018). Mesenchymal Stromal/Stem Cells in Regenerative Medicine and Tissue Engineering. Stem Cells Int 2018, 1–16.

Forstmeier, W., Segelbacher, G., Mueller, J.C., and Kempenaers, B. (2007). Genetic variation and differentiation in captive and wild zebra finches (Taeniopygia guttata). Molecular Ecology 16, 4039–4050.

Fritz, J.A., Brancale, J., Tokita, M., Burns, K.J., Hawkins, M.B., Abzhanov, A., and Brenner, M.P. (2014). Shared developmental programme strongly constrains beak shape diversity in songbirds. Nature Communications 5, 1–9.

Fu, Y., Chen, Z., Li, C., and Liu, G. (2012). Establishment of a duck cell line susceptible to duck hepatitis virus type 1. Journal of Virological Methods 184, 41–45.

Garcia-Oscos, F., Koch, T.M.I., Pancholi, H., Trusel, M., Daliparthi, V., Co, M., Park, S.E., Ayhan, F., Alam, D.H., Holdway, J.E., et al. (2021). Autism-linked gene FoxP1 selectively regulates the cultural transmission of learned vocalizations. Sci Adv 7, eabd2827.

Gedman, G.L., Wirthlin, M.L., Pfenning, A., and Jarvis, E. (2021). Convergent brain molecular specializations for vocal imitation in songbirds and humans. Manuscript in Preparation.

Gessara, I., Dittrich, F., Hertel, M., Hildebrand, S., Pfeifer, A., Frankl-Vilches, C., McGrew, M., and Gahr, M. (2021). Highly Efficient Genome Modification of Cultured Primordial Germ Cells with Lentiviral Vectors to Generate Transgenic Songbirds. Stem Cell Rep.

Griffiths, R., Double, M.C., Orr, K., and Dawson, R.J.G. (1998). A DNA test to sex most birds. Mol Ecol 7, 1071–1075.

Guan, D., McCarthy, S.A., Wood, J., Howe, K., Wang, Y., and Durbin, R. (2020). Identifying and removing haplotypic duplication in primary genome assemblies. Bioinformatics 36, 2896–2898.

Hafemeister, C., and Satija, R. (2019). Normalization and variance stabilization of single-cell RNA-seq data using regularized negative binomial regression. Genome Biology 20, 296–15.

Hara, E., Rivas, M.V., Ward, J.M., Okanoya, K., and Jarvis, E.D. (2012). Convergent Differential Regulation of Parvalbumin in the Brains of Vocal Learners. Plos One 7, e29457.

Heston, J.B., and White, S.A. (2017). To transduce a zebra finch: interrogating behavioral mechanisms in a model system for speech. Journal of Comparative Physiology A 203, 691–706.

Horita, H., Wada, K., Rivas, M.V., Hara, E., and Jarvis, E.D. (2010). The dusp1 immediate early gene is regulated by natural stimuli predominantly in sensory input neurons. J Comp Neurol 518, 2873–2901.

Hughes, P., Marshall, D., Reid, Y., Parkes, H., and Gelber, C. (2007). The costs of using unauthenticated, over-passaged cell lines: how much more data do we need? BioTechniques 43, 575–586.

Itoh, Y., and Arnold, A.P. (2011). Zebra finch cell lines from naturally occurring tumors. In Vitro Cellular & Developmental Biology. Animal 47, 280–282.

Itoh, Y., Kampf, K., Balakrishnan, C.N., and Arnold, A.P. (2011). Karyotypic polymorphism of the zebra finch Z chromosome. Chromosoma 120, 255–264.

Jarvis, E.D., Yu, J., Rivas, M.V., Horita, H., Feenders, G., Whitney, O., Jarvis, S.C., Jarvis, E.R., Kubikova, L., Puck, A.E.P., et al. (2013). Global view of the functional molecular organization of the avian cerebrum: Mirror images and functional columns. J Comp Neurol 521, 3614–3665.

Jarvis, E.D., Mirarab, S., Aberer, A.J., Li, B., Houde, P., Li, C., Ho, S.Y.W., Faircloth, B.C., Nabholz, B., Howard, J.T., et al. (2014). Whole-genome analyses resolve early branches in the tree of life of modern birds. Science 346, 1320–1331.

Jensen, K.T., Fløe, L., Petersen, T.S., Huang, J., Xu, F., Bolund, L., Luo, Y., and Lin, L. (2017). Chromatin accessibility and guide sequence secondary structure affect CRISPR‐Cas9 gene editing efficiency. Febs Lett 591, 1892–1901.

Joung, J., Konermann, S., Gootenberg, J.S., Abudayyeh, O.O., Platt, R.J., Brigham, M.D., Sanjana, N.E., and Zhang, F. (2017). Genome-scale CRISPR-Cas9 knockout and transcriptional activation screening. Nature Protocols 12, 828–863.

Jung, K.M., Kim, Y.M., Keyte, A.L., Biegler, M.T., Rengaraj, D., Lee, H.J., Mello, C.V., Velho, T.A.F., Fedrigo, O., Haase, B., et al. (2019). Identification and characterization of primordial germ cells in a vocal learning Neoaves species, the zebra finch. Faseb J 33, 13825–13836.

Kastenhuber, E.R., and Lowe, S.W. (2017). Putting p53 in Context. Cell 170, 1062–1078.

Katayama, M., Kiyono, T., Ohmaki, H., Eitsuka, T., Endoh, D., Inoue‐Murayama, M., Nakajima, N., Onuma, M., and Fukuda, T. (2019). Extended proliferation of chicken‐ and Okinawa rail‐derived fibroblasts by expression of cell cycle regulators. J Cell Physiol 234, 6709–6720.

Kinsella, C.M., Ruiz-Ruano, F.J., Dion-Côté, A.-M., Charles, A.J., Gossmann, T.I., Cabrero, J., Kappei, D., Hemmings, N., Simons, M.J.P., Camacho, J.P.M., et al. (2019). Programmed DNA elimination of germline development genes in songbirds. Nature Communications 10, 5468–10.

Köhler, S., Gargano, M., Matentzoglu, N., Carmody, L.C., Lewis-Smith, D., Vasilevsky, N.A., Danis, D., Balagura, G., Baynam, G., Brower, A.M., et al. (2020). The Human Phenotype Ontology in 2021. Nucleic Acids Res 49, gkaa1043..

Korlach, J., Gedman, G., Kingan, S.B., Chin, C.-S., Howard, J.T., Audet, J.-N., Cantin, L., and Jarvis, E.D. (2017). De novo PacBio long-read and phased avian genome assemblies correct and add to reference genes generated with intermediate and short reads. GigaScience 6, 695–16.

Kulak, M., Takki, O., and Galkina, S. (2020). Cell Culture Establishment from Zebra Finch Embryonic Fibroblasts. Adv Animal Vet Sci 9.

Laherty, C.D., Billin, A.N., Lavinsky, R.M., Yochum, G.S., Bush, A.C., Sun, J.-M., Mullen, T.-M., Davie, J.R., Rose, D.W., Glass, C.K., et al. (1998). SAP30, a Component of the mSin3 Corepressor Complex Involved in N-CoR-Mediated Repression by Specific Transcription Factors. Molecular Cell 2, 33–42.

Landry, G.P. (1997). An Almost Complete Guide to: The Varieties and Genetics of the Zebra Finch (Franklin, Louisiana, USA: Poule d’eau Publishing Co.).

Lane, D.P., and Crawford, L.V. (1979). T antigen is bound to a host protein in SV40-transformed cells. Nature 278, 261–263.

Li, H. (2018). Minimap2: pairwise alignment for nucleotide sequences. Bioinformatics 34, 3094–3100.

Li, H., Handsaker, B., Wysoker, A., Fennell, T., Ruan, J., Homer, N., Marth, G., Abecasis, G., Durbin, R., and Subgroup, 1000 Genome Project Data Processing (2009). The Sequence Alignment/Map format and SAMtools. Bioinformatics 25, 2078–2079.

Li, Z., Michael, I.P., Zhou, D., Nagy, A., and Rini, J.M. (2013). Simple piggyBac transposon-based mammalian cell expression system for inducible protein production. Proc National Acad Sci 110, 5004–5009.

Lin, Y.-C., Balakrishnan, C.N., and Clayton, D.F. (2014a). Functional genomic analysis and neuroanatomical localization of miR-2954, a song-responsive sex-linked microRNA in the zebra finch. Frontiers in Neuroscience 8, R106.

Lin, Y.-C., Boone, M., Meuris, L., Lemmens, I., Roy, N.V., Soete, A., Reumers, J., Moisse, M., Plaisance, S., Drmanac, R., et al. (2014b). Genome dynamics of the human embryonic kidney 293 lineage in response to cell biology manipulations. Nature Communications 5, 8.

Long, M.A., and Fee, M.S. (2008). Using temperature to analyse temporal dynamics in the songbird motor pathway. Nature 456, 189–194.

Lovell, P.V., Wirthlin, M., Kaser, T., Buckner, A.A., Carleton, J.B., Snider, B.R., McHugh, A.K., Tolpygo, A., Mitra, P.P., and Mello, C.V. (2020). ZEBrA: Zebra finch Expression Brain Atlas—A resource for comparative molecular neuroanatomy and brain evolution studies. J Comp Neurol 528, 2099–2131.

Macville, M., Schröck, E., Padilla-Nash, H., Keck, C., Ghadimi, B.M., Zimonjic, D., Popescu, N., and Ried, T. (1999). Comprehensive and Definitive Molecular Cytogenetic Characterization of HeLa Cells by Spectral Karyotyping. Cancer Research 59, 141–150.

Masters, J.R., and Stacey, G.N. (2007). Changing medium and passaging cell lines. Nature Protocols 2, 2276–2284.

McDowell, K.A., Hutchinson, A.N., Wong-Goodrich, S.J.E., Presby, M.M., Su, D., Rodriguiz, R.M., Law, K.C., Williams, C.L., Wetsel, W.C., and West, A.E. (2010). Reduced Cortical BDNF Expression and Aberrant Memory in Carf Knock-Out Mice. J Neurosci 30, 7453–7465.

Mello, C.V. (2014). The Zebra Finch, Taeniopygia guttata: An Avian Model for Investigating the Neurobiological Basis of Vocal Learning. Cold Spring Harbor Protocols 2014, 1237–1242.

Murray, J.R., Varian-Ramos, C.W., Welch, Z.S., and Saha, M.S. (2013). Embryological staging of the Zebra Finch, Taeniopygia guttata. Journal of Morphology 274, 1090–1110.

Nelson, C.E., Hakim, C.H., Ousterout, D.G., Thakore, P.I., Moreb, E.A., Rivera, R.M.C., Madhavan, S., Pan, X., Ran, F.A., Yan, W.X., et al. (2015). In vivo genome editing improves muscle function in a mouse model of Duchenne muscular dystrophy. Science aad5143.

Newhouse, D.J., Hofmeister, E.K., and Balakrishnan, C.N. (2017). Transcriptional response to West Nile virus infection in the zebra finch (Taeniopygia guttata). Royal Society Open Science 4, 170296.

Nihashi, Y., Umezawa, K., Shinji, S., Hamaguchi, Y., Kobayashi, H., Kono, T., Ono, T., Kagami, H., and Takaya, T. (2019). Distinct cell proliferation, myogenic differentiation, and gene expression in skeletal muscle myoblasts of layer and broiler chickens. Sci Rep-Uk 9, 16527.

O’Brien, A., and Bailey, T.L. (2014). GT-Scan: identifying unique genomic targets. Bioinformatics 30, 2673–2675.

Olson, C.R., Hodges, L.K., and Mello, C.V. (2015). Dynamic gene expression in the song system of zebra finches during the song learning period. Developmental Neurobiology n/a-n/a.

Petkov, C.I., and Jarvis, E. (2012). Birds, primates, and spoken language origins: behavioral phenotypes and neurobiological substrates. Frontiers in Evolutionary Neuroscience 4.

Pfenning, A.R., Hara, E., Whitney, O., Rivas, M.V., Wang, R., Roulhac, P.L., Howard, J.T., Wirthlin, M., Lovell, P.V., Ganapathy, G., et al. (2014). Convergent transcriptional specializations in the brains of humans and song-learning birds. Science 346, 1256846–1256846.

Pigozzi, M.I., and Solari, A.J. (1998). Germ cell restriction and regular transmission of an accessory chromosome that mimics a sex body in the zebra finch, Taeniopygia guttata. Chromosome Res 6, 105–113.

Pinello, L., Canver, M.C., Hoban, M.D., Orkin, S.H., Kohn, D.B., Bauer, D.E., and Yuan, G.-C. (2016). Analyzing CRISPR genome-editing experiments with CRISPResso. Nature Biotechnology 34, 695–697.

Quinlan, A.R., and Hall, I.M. (2010). BEDTools: a flexible suite of utilities for comparing genomic features. Bioinformatics 26, 841–842.

Ran, F.A., Cong, L., Yan, W.X., Scott, D.A., Gootenberg, J.S., Kriz, A.J., Zetsche, B., Shalem, O., Wu, X., Makarova, K.S., et al. (2015). In vivo genome editing using Staphylococcus aureus Cas9. Nature 520, 186–191.

Ray, F.A., Meyne, J., and Kraemer, P.M. (1992). SV40 T antigen induced chromosomal changes reflect a process that is both clastogenic and aneuploidogenic and is ongoing throughout neoplastic progression of human fibroblasts. Mutat Res Fundam Mol Mech Mutagen 284, 265–273.

Reyes, R.C., Brennan, A.M., Shen, Y., Baldwin, Y., and Swanson, R.A. (2012). Activation of Neuronal NMDA Receptors Induces Superoxide-Mediated Oxidative Stress in Neighboring Neurons and Astrocytes. J Neurosci 32, 12973–12978.

Rhie, A., McCarthy, S.A., Fedrigo, O., Damas, J., Formenti, G., Koren, S., Uliano-Silva, M., Chow, W., Fungtammasan, A., Gedman, G.L., et al. (2020). Towards complete and error-free genome assemblies of all vertebrate species. BioRxiv 3, 2020.05.22.110833.

Rhie, A., McCarthy, S.A., Fedrigo, O., Damas, J., Formenti, G., Koren, S., Uliano-Silva, M., Chow, W., Fungtammasan, A., Kim, J., et al. (2021). Towards complete and error-free genome assemblies of all vertebrate species. Nature 592, 737–746.

Robinson, J.T., Thorvaldsdóttir, H., Winckler, W., Guttman, M., Lander, E.S., Getz, G., and Mesirov, J.P. (2011). Integrative genomics viewer. Nat Biotechnol 29, 24–26.

Rosselló, R.A., Chen, C.-C., Dai, R., Howard, J.T., Hochgeschwender, U., and Jarvis, E.D. (2013). Mammalian genes induce partially reprogrammed pluripotent stem cells in non-mammalian vertebrate and invertebrate species. Elife 2, e00036.

Rossello, R.A., Pfenning, A., Howard, J.T., and Hochgeschwender, U. (2016). Characterization and genetic manipulation of primed stem cells into a functional naïve state with ESRRB. World J Stem Cells 8, 355.

Santos, M. da S. dos, Kretschmer, R., Frankl-Vilches, C., Bakker, A., Gahr, M., O’Brien, P.C.M., Ferguson-Smith, M.A., and Oliveira, E.H.C. de (2017). Comparative Cytogenetics between Two Important Songbird, Models: The Zebra Finch and the Canary. Plos One 12, e0170997.

Schaefer-Klein, J., Givol, I., Barsov, E.V., Whitcomb, J.M., VanBrocklin, M., Foster, D.N., Federspiel, M.J., and Hughes, S.H. (1998). The EV-O-Derived Cell Line DF-1 Supports the Efficient Replication of Avian Leukosis-Sarcoma Viruses and Vectors. Virology 248, 305–311.

Shah, Z.H., Jones, D.R., Sommer, L., Foulger, R., Bultsma, Y., D’Santos, C., and Divecha, N. (2013). Nuclear phosphoinositides and their impact on nuclear functions. FEBS Journal 280, 6295–6310.

Steyaert, S., Diddens, J., Galle, J., Meester, E.D., Keulenaer, S.D., Bakker, A., Sohnius-Wilhelmi, N., Frankl-Vilches, C., Linden, A.V. der, Criekinge, W.V., et al. (2016). A genome-wide search for eigenetically regulated genes in zebra finch using MethylCap-seq and RNA-seq. Scientific Reports 6, 20957.

Velho, T.A.F., Lovell, P.V., Friedrich, S.R., Olson, C.R., Miles, J., Mueller, P.A., Tavori, H., Fazio, S., Lois, C., and Mello, C.V. (2021). Divergent low-density lipoprotein receptor (LDLR) linked to low VSV G-dependent viral infectivity and unique serum lipid profile in zebra finches. Proc National Acad Sci 118, e2025167118.

Viiri, K.M., Korkeamäki, H., Kukkonen, M.K., Nieminen, L.K., Lindfors, K., Peterson, P., Mäki, M., Kainulainen, H., and Lohi, O. (2006). SAP30L interacts with members of the Sin3A corepressor complex and targets Sin3A to the nucleolus. Nucleic Acids Res 34, 3288–3298.

Viiri, K.M., Jänis, J., Siggers, T., Heinonen, T.Y.K., Valjakka, J., Bulyk, M.L., Mäki, M., and Lohi, O. (2009). DNA-binding and -bending activities of SAP30L and SAP30 are mediated by a zinc-dependent module and monophosphoinositides. Molecular and Cellular Biology 29, 342–356.

Vunjak-Novakovic, G., and Freshney, R.I. (2006). Culture of Cells for Tissue Engineering (John Wiley & Sons).

Wade, J., and Arnold, A.P. (2004). Sexual differentiation of the zebra finch song system. Annals of the New York Academy of Sciences 1016, 540–559.

Warren, W.C., Clayton, D.F., Ellegren, H., Arnold, A.P., Hillier, L.W., Künstner, A., Searle, S., White, S., Vilella, A.J., Fairley, S., et al. (2010). The genome of a songbird. Nature 464, 757–762.

Watanabe, Y., Kameoka, S., Gopalakrishnan, V., Aldape, K.D., Pan, Z.Z., Lang, F.F., and Majumder, S. (2004). Conversion of myoblasts to physiologically active neuronal phenotype. Gene Dev 18, 889–900.

Winding, P., and Berchtold, M.W. (2001). The chicken B cell line DT40: a novel tool for gene disruption experiments. Journal of Immunological Methods 249, 1–16.

Wirthlin, M., Lovell, P.V., Jarvis, E.D., and Mello, C.V. (2014). Comparative genomics reveals molecular features unique to the songbird lineage. Bmc Genomics 15, 1082.

Xiao, D., Liu, X., Zhang, M., Zou, M., Deng, Q., Sun, D., Bian, X., Cai, Y., Guo, Y., Liu, S., et al. (2018). Direct reprogramming of fibroblasts into neural stem cells by single non-neural progenitor transcription factor Ptf1a. Nature Communications 9, 1–19.

Yang, C., Zhou, Y., Marcus, S., Formenti, G., Bergeron, L.A., Song, Z., Bi, X., Bergman, J., Rousselle, M.M.C., Zhou, C., et al. (2021). Evolutionary and biomedical insights from a marmoset diploid genome assembly. Nature 1–7.

Zhang, Y., Sun, Z.-W., Iratni, R., Erdjument-Bromage, H., Tempst, P., Hampsey, M., and Reinberg, D. (1998). SAP30, a Novel Protein Conserved between Human and Yeast, Is a Component of a Histone Deacetylase Complex. Molecular Cell 1, 1021–1031.

Zhou, X., Hollern, D., Liao, J., Andrechek, E., and Wang, H. (2013). NMDA receptor-mediated excitotoxicity depends on the coactivation of synaptic and extrasynaptic receptors. Cell Death Dis 4, e560–e560.

